# Mechanistic basis for SNX27-Retromer coupling to ESCPE-1 in promoting endosomal cargo recycling

**DOI:** 10.1101/2021.08.28.457928

**Authors:** Boris Simonetti, Qian Guo, Manuel Gimenez-Andres, Kai-En Chen, Edmund R.R. Moody, Ashley J. Evans, Chris M. Danson, Tom A. Williams, Brett M. Collins, Peter J. Cullen

## Abstract

Sorting nexin-27 (SNX27)-Retromer is an endosomal sorting complex that orchestrates endosome-to-plasma membrane recycling of hundreds of internalized receptors, channels and transporters, enzymes and adhesion molecules. While SNX27-Retromer is essential for development, subtle functional defects are observed in human disease, most notably neurodegenerative and neurological disorders. Achieving a thorough mechanistic dissection of SNX27-Retromer is central to understanding endosomal sorting in health and disease. Here we combine biochemical, structural and cellular analyses to establish the mechanistic basis through which SNX27-Retromer couples to the membrane tubulating ESCPE-1 complex (Endosomal SNX-BAR sorting complex for promoting exit 1). We show that a conserved surface in the FERM (4.1/ezrin/radixin/moesin) domain of SNX27 directly binds acidic-Asp-Leu-Phe (aDLF) motifs in the disordered amino-termini of the SNX1 and SNX2 subunits of ESCPE-1. This interaction hands-over SNX27-Retromer captured integral membrane proteins into ESCPE-1 tubular profiles to promote their cell surface recycling. Through phylogenetic analysis, we reveal that SNX27:Retromer:ESCPE-1 assembly evolved in a stepwise manner during the early evolution of metazoans, which reflects the increasing complexity of endosomal sorting from the ancestral opisthokont to modern animals.

## INTRODUCTION

A central role of the endosomal-lysosomal network is to determine the sorting of integral proteins and their associated proteins and lipids, collectively termed ‘cargos’, between two fates, either degradation within lysosomes or retrieval from this fate for the promotion of recycling to a variety of organelles that include the cell surface (Cullen and Steinberg, 2018). Efficient and selective endosomal-lysosomal cargo sorting is essential for the function of all eukaryotic cells with increasing evidence pointing to defects in endosomal retrieval and recycling being at the heart of a wide array of human diseases most notably those associated with neurodegeneration (Small et al., 2020). Establishing the molecular mechanisms and functional significance of endosomal cargo retrieval and recycling is therefore fundamental to understand eukaryotic cell biology.

Two general concepts are considered to account for the mechanism of endosomal cargo retrieval and recycling. Sequence-independent sorting posits that sorting can proceed without the need for any sequence information within the intracellular cytoplasmic domains of integral membrane cargos (Maxfield and McGraw, 2004). On the other hand, sequence-dependent sorting is controlled by specific signals in the cytoplasmic domain of cargos. These signals are typically short linear peptide motifs that direct the sorting process through coupling to ancient and evolutionary conserved coat complexes (Cullen and Steinberg, 2018; McNally and Cullen, 2018; Chen et al., 2019). In coupling sorting motif recognition with membrane re-modelling, these coats co-ordinate the process of cargo retrieval with the formation of cargo-enriched transport carriers for onward recycling. In mammals, these coats include sorting nexin-3 (SNX3)-Retromer (Strochlic et al., 2007; Harterink et al., 2011; Lucas et al., 2016; Kendall et al., 2020; Leneva et al., 2021), SNX27-Retromer (Lauffer et al., 2010; Temkin et al., 2011; Steinberg et al., 2013; Gallon et al., 2014), SNX17-Retriever (McNally et al., 2017), the CCC (CCDC22/CCDC93/COMMD) (Phillips-Krawczak et al., 2015; Bartuzi et al., 2016) and ESCPE-1 (Endosomal SNX-BAR sorting complex for promoting exit 1) complexes (Simonetti et al., 2019; Yong et al., 2020), and various accessory proteins including the actin polymerizing WASH (Wiskott–Aldrich syndrome protein and SCAR homologue) complex (Gomez and Billadeau, 2009; Derivery et al., 2009).

Sorting nexin-27 (SNX27) is an endosome-associated cargo adaptor composed of a central PX (phox homology) domain flanked by an N-terminal PDZ (PSD95, discs large, zona occludens) domain and a C-terminal FERM (4.1/ezrin/radixin/moesin) domain (Kajii et al., 2003; Ghai et al., 2011). The PDZ domain recognizes a specific sorting motif, termed the PDZ binding motif (PDZbm), present at the very C-termini of over 400 cargo proteins including receptors, transporters, channels, enzymes and adhesion molecules (Joubert et al., 2004; Lunn et al., 2007; Lauffer et al., 2010; Balana et al., 2011; Cai et al., 2011; Ghai et al., 2011; Temkin et al., 2011; Steinberg et al., 2013; Hussain et al., 2014; Clairfeuille et al., 2016; Sharma et al., 2020; McMillan et al., 2021). In recognizing these internalized cargos, SNX27 regulates their retrieval from lysosomal degradation and promotes their recycling back to the plasma membrane (Lauffer et al., 2010; Temkin et al., 2011; Steinberg et al., 2013).

To facilitate the retrieval process, the SNX27 PDZ domain associates with Retromer, a stable heterotrimer of VPS26 (A and B isoforms expressed in humans (Kerr et al., 2005)), VPS35 and VPS29, to form the SNX27-Retromer (Temkin et al., 2011; Steinberg et al., 2013; Gallon et al., 2014). Retromer itself assembles to form dimeric arches that may cluster SNX27 captured cargos (Kovtun et al., 2018; Kendall et al., 2020; Leneva et al., 2021) and associates with the WASH complex to allow the localized Arp2/3-dependent polymerisation of branched filamentous actin (Harbour et al., 2012; Jia et al., 2012). In a poorly understood process, these assemblies and actin dynamics are proposed to generate a retrieval sub-domain on the endosomal membrane that is enriched in cargo destined for recycling (Puthenveedu et al., 2010; Jia et al., 2012; Tsvetanova et al., 2015; Varandas et al., 2016; Lee et al., 2016). It is from these sub-domains that further membrane re-modelling ensues to drive the biogenesis of cargo-enriched tubular profiles and tubulovesicular carriers that physically transport cargo back to the cell surface.

Importantly, defects in SNX27-Retromer function are associated with the pathoetiology of human disease most notable neurological disorders and neurodegenerative disease (Chandra et al., 2021).

A membrane tubulating BAR (Bim/Amphiphysin/Rvs) domain-containing protein associated with SNX27-Retromer tubular profiles is sorting nexin-1 (SNX1) (Temkin et al., 2011). SNX1, and its functional ortholog SNX2, are part of the ESCPE-1 complex (Carlton et al., 2004; Simonetti et al., 2019; Yong et al., 2020), a membrane tubulating complex that *in vitro* can reconstitute the formation of membrane tubules with diameters similar to those observed *in cellulo* (van Weering et al., 2012). To be functional, ESCPE-1 forms a stable heterodimer of either SNX1 or SNX2 associated with either of SNX5 or SNX6, two additional BAR domain-containing orthologs (Wassmer et al., 2007). Importantly, the mechanistic basis of how SNX27-Retromer delivers cargo into forming ESCPE-1 tubular profiles and tubulovesicular carriers remains a major unanswered question.

Here we establish that a conserved surface in the FERM domain of SNX27 binds directly to acidic-Asp-Leu-Phe (aDLF) sequence motifs present in the disordered N-termini of the SNX1 and SNX2 subunits of ESCPE-1. We show that this interaction is required to hand-over PDZ-binding motif containing cargo proteins retrieved from lysosomal degradation by SNX27-Retromer into ESCPE-1 tubular profiles that promote cargo recycling for subsequent repopulation at the cell surface. Through evolutionary analysis of those protein-protein interactions that assemble the SNX27:Retromer:ESCPE-1 coat complex, we reveal how this sorting axis has evolved to accommodate the increasing complexities of endosomal sorting observed in animals.

## RESULTS

### The FERM domain of SNX27 promotes cargo recycling following SNX27-Retromer mediated lysosomal retrieval

Previously, we have established that the interaction of Retromer with the PDZ domain of SNX27 is a crucial step in preventing the routing of internalised cargo into the lysosomal degradative pathway (Steinberg et al., 2013; Gallon et al., 2014). To identify the region(s) of SNX27 required for promoting subsequent cargo recycling to the plasma membrane we used CRISPR gene-editing to generate SNX27 knock-out HeLa cells (Figure 1A). Compared to control cells, where endogenous GLUT1 was enriched at the cell surface, in the SNX27 knock-out cells GLUT1 displayed a steady-state enrichment with LAMP1-positive lysosomes, a phenotype that was also observed in a VPS35 knock-out HeLa cell line (Figure 1A). Importantly, re-expression of GFP-SNX27 in the SNX27 knock-out cells rescued the retrieval of GLUT1 from the lysosomal distribution and promoted its repopulation at the cell surface (Figure 1B). These data mirror those previously published using RNAi-mediated SNX27 suppression (Steinberg et al., 2013), and confirm the SNX27 knock-out HeLa cell line as a suitable screening platform through which to dissect the features of SNX27 required for promoting GLUT1 endosome-to-plasma membrane recycling.

**Figure 1.**
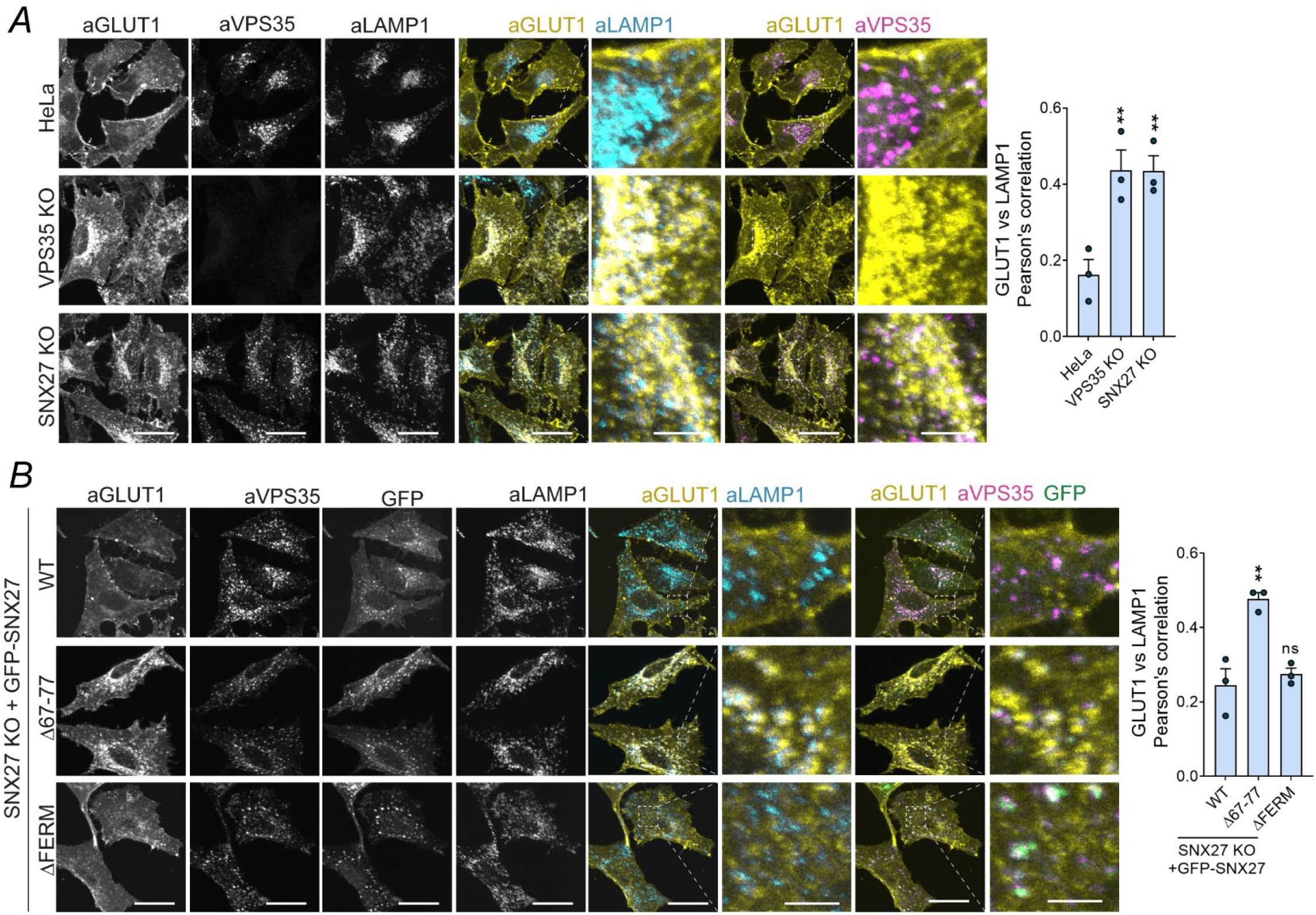
The FERM domain of SNX27 promotes cargo recycling following SNX27-Retromer mediated lysosomal retrieval. **A** Representative images of parental Hela cells, VPS35 KO HeLa clonal line and SNX27 KO HeLa cells. The steady-state distribution of GLUT1 was analyzed by immunofluorescence staining of fixed cells with GLUT1 and the endosomal and lysosomal markers VPS35 and LAMP1. 50 cells were analyzed in each condition for colocalization between GLUT1 and LAMP1 across n = 3 independent experiments. The Pearson’s coefficient values were compared to the values of parental HeLa using one-way ANOVA and Dunnett’s test: VPS35 KO vs HeLa, P = 0.0089; SNX27 KO vs HeLa, P = 0.0093. **B** Representative images of SNX27 KO HeLa cells transiently transfected with GFP-SNX27 WT, GFP-SNX27 Δ67-77 and GFP-SNX27 ΔFERM. 48h after transfection, cells were fixed and immunostained. The steady-state distribution of GLUT1 was analyzed by immunofluorescence staining of fixed cells with GLUT1 and the endosomal and lysosomal markers VPS35 and LAMP1. 50 cells were analyzed in each condition for colocalization between GLUT1 and LAMP1 across n = 3 independent experiments. The Pearson’s coefficient values were compared to the values of SNX27 KO cells rescued with GFP-SNX27 WT using one-way ANOVA and Dunnett’s test: GFP-SNX27Δ67-77 vs GFP-SNX27 WT, P = 0.0024; GFP-SNX27ΔFERM KO vs GFP-SNX27 WT, P = 0.7008. Bars, error bars and symbols represent the mean, s.e.m. and individual data points, respectively. *P< 0.05, **P< 0.01, ***P< 0.001, ****P< 0.0001, ns = not significant. Scale bars, 25 μm (micrographs) and 5 μm (magnified images).

In contrast, expression of wild-type SNX27, expression of a SNX27 mutant lacking the loop region essential for Retromer association (GFP-SNX27Δ67-77) (Steinberg et al., 2013; Gallon et al., 2014), failed to rescue lysosomal retrieval and failed to promote GLUT1 cell surface recycling (Figure 1B). These data further confirm that the association of SNX27 with Retromer is a crucial step in coupling sequence-dependent cargo recognition with cargo retrieval from the lysosomal degradative fate (Steinberg et al., 2013; Gallon et al., 2014). Consistent with this, expression of a SNX27 deletion mutant lacking the FERM domain (GFP-SNX27ΔFERM), a mutant that retained the association of SNX27 with Retromer, retained the ability to retrieve GLUT1 from lysosomes, as defined by a decreased colocalization between GLUT1 and LAMP1 (Figure 1B). However, the GFP-SNX27ΔFERM mutant failed to promote GLUT1 recycling to the cell surface, rather GLUT1 was enriched in a Retromer-positive endosomal compartment (Figure 1B). Together these data are consistent with a model in which the coupling of SNX27 to Retromer is necessary for retrieval from the lysosomal degradative fate while the SNX27 FERM domain is required to promote the subsequent endosome-to-cell surface recycling of GLUT1.

### SNX1 and SNX2 couple ESCPE-1 to the FERM domain of SNX27

ESCPE-1 regulates tubular-based endosomal recycling of several cargos, many of which are also dependent on SNX27-Retromer (Steinberg et al., 2013; Simonetti et al., 2019; Yong et al., 2020). ESCPE-1 is a heterodimeric assembly of the functionally redundant SNX1 or SNX2 associated with the functionally redundant SNX5 or SNX6 (Figure 2A) (Simonetti et al., 2019; Yong et al., 2020), while SNX27-Retromer is an assembly of SNX27 with the VPS26:VPS35:VPS29 heterotrimeric Retromer complex (Figure 2B) (Steinberg et al., 2013). To define a possible connection between SNX27 and ESCPE-1 we first performed a series of GFP-nanotrap immuno-isolation of SNX27 and various SNX27 mutants expressed in HEK-293T cells and probed for the presence of associated endogenous ESCPE-1. Robust association was observed between full length GFP-SNX27 and all components of ESCPE-1 (Figure 2C, 2D and 2E). This association was independent of the SNX27 PDZ domain as binding was retained in the PDZ deletion mutant GFP-SNX27ΔPDZ and was not observed when just the PDZ domain was expressed in HEK-293T cells (Figure 2C, 2D). Retromer association with SNX27 was not required for ESCPE-1 binding as GFP-SNX27Δ67-77 and GFP-SNX27(L67A/L77A), two mutants that fail to bind to Retromer (Gallon et al., 2014), each retained association with ESCPE-1 (Figure 2E). However, ESCPE-1 association was lost upon deletion of the SNX27 FERM domain (SNX27ΔFERM) or upon switching the FERM domain of SNX27 for the corresponding FERM domain from SNX17 (Figure 2C). Moreover, just the isolated FERM domain of SNX27, when expressed in HEK-293T cells, was able to associate with ESCPE-1 (Figure 2D). Finally, in the reverse immuno-isolations, GFP-tagged SNX1, SNX2, SNX5, or SNX6 revealed that only SNX1 and SNX2 displayed a pronounced association with endogenous SNX27 when expressed in HEK-293T cells (Figure 2F). Moreover, this association did not require the presence of the SNX1 or SNX2 BAR domains, since deletion of these domains failed to affect binding to SNX27 (Figure 2G). Together these data are consistent with the FERM domain of SNX27 associating with the SNX1 and SNX2 components of ESCPE-1.

**Figure 2.**
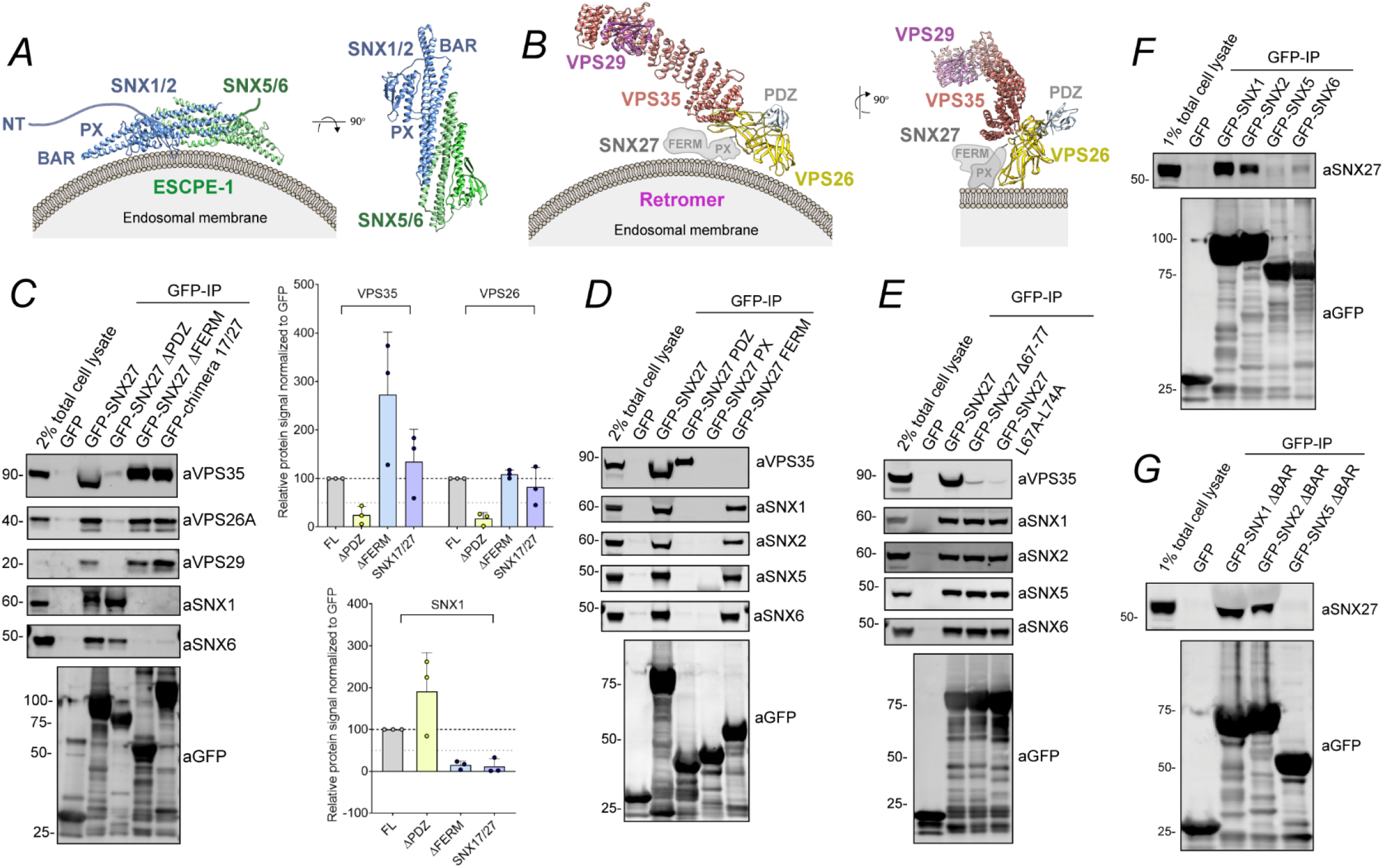
The FERM domain of SNX27 interacts with SNX1 and SNX2. **A** Schematics of ESCPE-1 assembly consisting of heterodimers of the SNX-BAR proteins SNX1 or SXN2 with SNX5 or SNX6. These SNXs have BAR and PX domains responsible of dimer formation and membrane targeting respectively. SNX1 and SNX2 also have an extended N-terminal unstructured extension (NT). **B** Schematics of the SNX27-Retromer assembly. Retromer is a stable heterotrimer of VPS35, VPS26 and VPS29. Retromer directly interacts with SNX27 through a β-hairpin insertion within the PDZ domain (residues 67-79) that engages a cleft of the VPS26 protein between the N- and C-terminal subdomains. **C** Co-immunoprecipitation of GFP-tagged SNX27 full length (FL), SNX27 ΔPDZ, SNX27 ΔFERM and a chimera consisting of the SNX27 gene where its FERM domain was swapped for that of SNX17. The constructs were transiently transfected in HEK-293T cells, the cell lysates were subjected to GFP-trap based immunoprecipitation and the immunoprecipitates were analyzed by quantitative fluorescence-based western blotting. The band intensities of VPS35, VPS26A, VPS29, SNX1, and GFP were measured from n= 3 independent experiments using Odyssey software. The band intensities, normalized to GFP expression, are presented as the average fraction of the full length. **D** Co-immunoprecipitation of GFP-tagged SNX27 full length, isolated SNX27 PDZ, PX and FERM domains expressed in HEK-293T cells. The cell lysates were subjected to GFP-trap based immunoprecipitation and the immunoprecipitates were blotted for VPS35, SNX1, SNX2, SNX5, SNX6 and GFP. The blot is representative of three independent GFP traps. **E** Co-immunoprecipitation of GFP-tagged SNX27 full length, and the SNX27 mutants Δ67-77 and L67A-L74A expressed in HEK-293T cells. The cell lysates were subjected to GFP-trap based immunoprecipitation and the immunoprecipitates were blotted for VPS35, SNX1, SNX2, SNX5, SNX6 and GFP. The blot is representative of three independent GFP traps. **F** Co-immunoprecipitation of GFP-tagged SNX1, SNX2, SNX5 and SNX6 expressed in HEK-293T cells. The cell lysates were subjected to GFP-trap based immunoprecipitation and the immunoprecipitates were blotted for SNX27 and GFP. The blot is representative of three independent GFP traps. **G** Co-immunoprecipitation of GFP-tagged SNX1, SNX2, SNX5 lacking the BAR domain (ΔBAR) expressed in HEK-293T cells. The cell lysates were subjected to GFP-trap based immunoprecipitation and the immunoprecipitates were blotted for SNX27 and GFP. The blot is representative of three independent GFP traps. Molecular masses are given in kilodaltons. Bars, error bars and symbols represent the mean, s.e.m. and individual data points, respectively.

### The N-termini of SNX1 and SNX2 present tandem acidic-Asp-Leu-Phe motifs that associate with the SNX27 FERM domain

Compared to SNX5 and SNX6, human SNX1 and SNX2 each contain N-terminal extensions of 142 and 139 amino acids respectively that are predicted to be unstructured. Expression in HEK-293T cells of GFP-SNX1(1-279), a construct encoding for the N-terminal extension plus the SNX1 PX domain, and GFP-SNX1(1-138), a construct encoding for just the N-terminal extension of SNX1, established that the N-terminal extension of SNX1 is sufficient for association with endogenous SNX27 (Figure 3A). To establish the direct nature of the association, we recombinantly expressed and purified full-length mouse SNX1 and the isolated FERM domain of human SNX27; mouse and human SNX1 are 94.2% identical. Isothermal titration calorimetry (ITC) established that SNX1 bound directly to the FERM domain of SNX27 with an affinity (*K*_d_) of 41.0 µM (Figure 3B; Supplementary Table 1). A similar binding affinity (39.1 µM) was observed with a recombinant protein corresponding to the N-terminal extension of mouse SNX1 (85.9% sequence identity with human SNX1) (Figure 3B). To map down the features of SNX1 and SNX2 required for SNX27 binding, we performed a deletion mutagenesis screen of the N-terminal extension of human SNX1 (Figure 3C). Walking upstream from a minimal binding region encoded by residues 1-138 established that two regions were required for maximal SNX27 binding. Deletion of residues 79 to 100 led to a pronounced reduction in binding and subsequent deletion of residues 41 to 54 abolished the interaction. Alignment of these regions of SNX1 revealed two similar motifs with DIF or DLF sequences flanked by upstream acidic residues. Comparison with the N-terminus of SNX2, which also binds SNX27 (Supplementary Figure 1A), showed that it also has two similar sequences in its disordered N-terminal domains, and altogether these define a conserved sequence D/E-x-x-D-I/L-F, here termed the acidic-D-L-F (aDLF) motif (Figure 3D and 3E).

**Figure 3.**
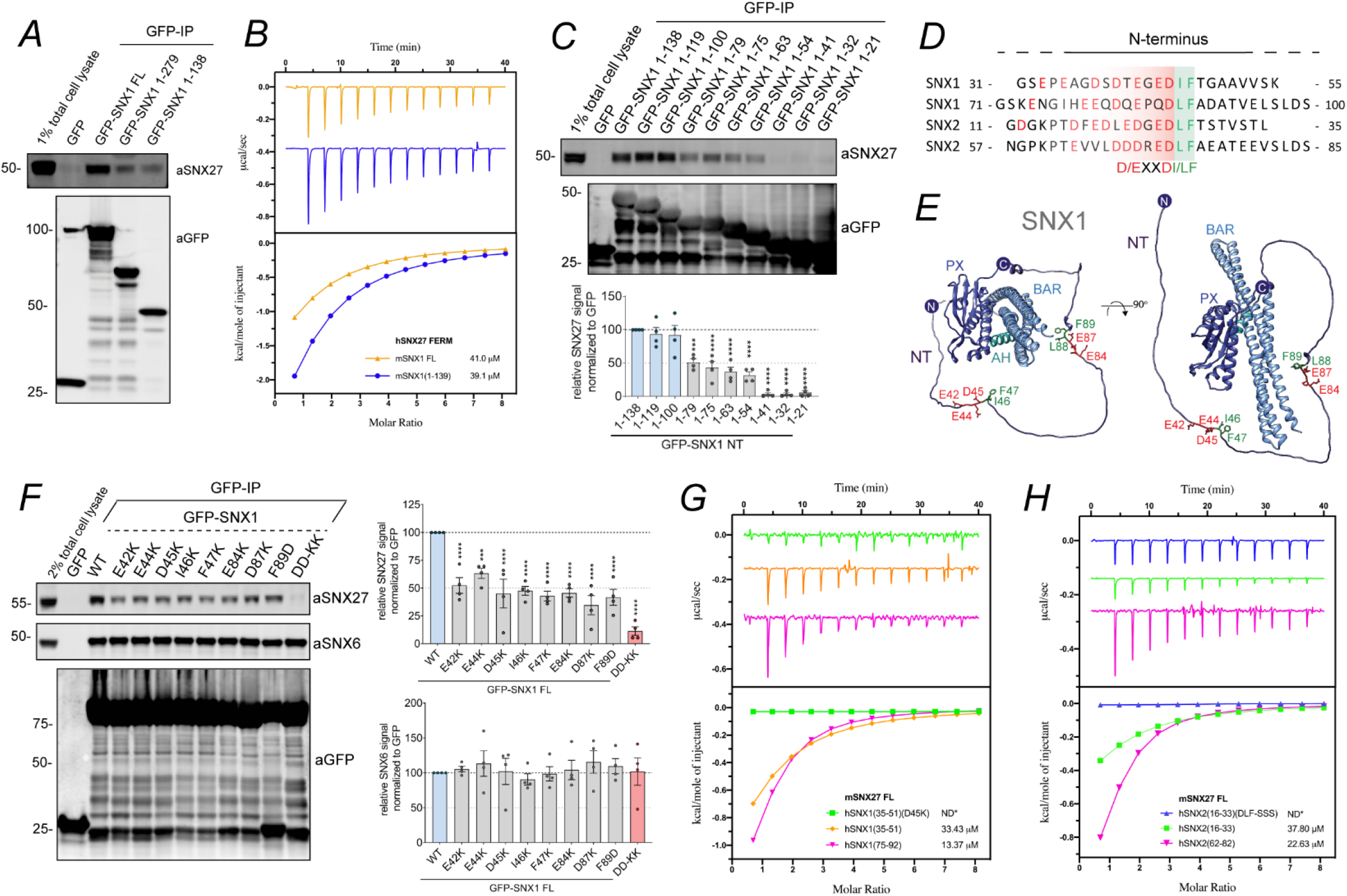
The N-termini of SNX1 and SNX2 directly interact with SNX27 FERM domain through tandem aDLF motifs. **A** Co-immunoprecipitation of GFP-tagged SNX1 full length (FL), or truncation mutants corresponding to residues 1-279 and 1-138 expressed in HEK-293T cells. The cell lysates were subjected to GFP-trap based immunoprecipitation and the immunoprecipitates were blotted for SNX27 and GFP. The blot is representative of three independent GFP traps. **B** Binding of mSNX1_FL_ (orange) and mSNX1_1-139_ (blue) with hSNX27_FERM_ by ITC. **C** Co-immunoprecipitation of GFP-tagged SNX1 N-terminus (1-138) and a panel of constructs corresponding to serial truncations of SNX1 N-terminus. The constructs were transiently transfected in HEK-293T cells, the cell lysates were subjected to GFP-trap based immunoprecipitation and the immunoprecipitates were analysed by quantitative fluorescence-based western blotting. The band intensity of SNX27 and GFP was measured from n= 4 independent experiments using Odyssey software and was normalized to GFP expression. The relative binding of SNX1 truncation mutants was compared with the SNX1 N-terminus (1-138) using a one-way ANOVA and Dunnett’s test. **D** Alignment of the SNX1 regions that are involved in SNX27 binding with the corresponding sequences in the SNX2 N-terminus. The conserved residues between SNX1 and SNX2 account for a consensus motif of acidic residues followed by DI/LF (aDLF motif). **E** Alphafold structural prediction for full length SNX1 (Tunyasuyunakool et al., 2021). The N-terminus of SNX1 is shown in its full extension and the residues involved in SNX27 binding are coloured in red and green. **F** Co-immunoprecipitation of GFP-tagged SNX1 wild type (WT) and a panel of amino acid swap mutants across the aDLF motifs of SNX1. The constructs were transiently transfected in HEK293T cells, the cell lysates were subjected to GFP-trap based immunoprecipitation and the immunoprecipitates were analysed by quantitative fluorescence-based western blotting. The band intensities of SNX27, SNX6 and GFP were measured from n= 4 independent experiments using Odyssey software. The band intensity, normalized to GFP expression, is presented as the average fraction of SNX1 WT. The binding of SNX1 point mutants to SNX27 and SNX6 were compared with the SNX1 WT using a one-way ANOVA and Dunnett’s test. **G** Binding of mSNX27_FL_ and hSNX1 peptides by ITC. SNX1_35-51(D45K)_: green; SNX1_35-51_: orange; hSNX1_75-92_: lilac; ND*: no binding detected. **H** Binding of mSNX27_FL_ and hSNX2 peptides by ITC. SNX2_16-33(DLF/SSS)_: blue; SNX2_16-33_: green; SNX2_62-82_: lilac; ND*: no binding detected. The ITC graphs represent the integrated and normalized data fit with 1:1 ratio binding. The binding affinity (*K*_d_) is given as mean of at least two independent experiments. Thermodynamic parameters for the binding are provided in Supplementary Table 1. Molecular masses are given in kilodaltons. Bars, error bars and symbols represent the mean, s.e.m. and individual data points, respectively. *P< 0.05, **P< 0.01, ***P< 0.001, ****P< 0.0001

Site-directed mutagenesis targeting the first aDLF motif of SNX1 established that full-length GFP-SNX1(E42K), GFP-SNX1(E44K), GFP-SNX1(D45K), GFP-SNX1(I46K) and GFP-SNX1(F47K) all displayed reduced binding to SNX27 of around 50% (Figure 3F). A similar reduction in binding was also observed when targeting the second aDLF motif of SNX1 as observed with the GFP-SNX1(E84K), GFP-SNX1(D87K) and GFP-SNX1(F89D) mutants (Figure 3F). In a double GFP-SNX1(D45K,D87K) mutant that simultaneously targeted key residues in both aDLF motifs SNX27 binding was reduced to near undetectable levels (Figure 3F). Importantly, all mutants retained their ability to associate with SNX6, establishing that ESCPE-1 was still able to assemble. Next, we used ITC to quantify the binding of aDLF peptides from SNX1 and SNX2 to full-length mouse SNX27. Peptides corresponding to residues 35 to 51 and 75 to 92 encompassing the first and second aDLF motifs of human SNX1 respectively, bound to SNX27 with respective affinities of 33.4 µM and 13.3 µM (Figure 3G; Supplementary Table 1). Establishing the redundant nature of the interaction with SNX27, we designed peptides corresponding to the first and second aDLF motifs of SNX2, namely residues 16 to 33 and 62 to 82. These also bound to full-length mouse SNX27 with affinities of 37.8 µM and 22.6 µM respectively (Figure 3H). Further defining the redundancy between SNX1 and SNX2, introduction of a charge swap mutation into the SNX1 35-51 peptide, SNX1(D45K), and a triple mutation in the SNX2 16-33 peptide, SNX2(DLF-SSS), resulted in mutant peptides that showed no detectable binding to SNX27 (Figure 3G and 3H). Finally, a competitive ITC assay using full-length SNX27 pre-incubated with excess SNX2 16 to 33 peptide established that it competed for binding of SNX1 75 to 92, consistent with both sequences associating with the same surface within the SNX27 FERM domain (Supplementary Figure 1B). Overall, these data establish that through tandem aDLF motifs present within their unstructured amino terminal extensions SNX1 and SNX2 directly associate with the FERM domain of SNX27.

### A basic surface in the SNX27 FERM domain is required for SNX1/SNX2 binding

SNX27 and SNX17 are related endosome associated cargo adaptors (Ghai et al., 2011; Ghai et al., 2013). While they display entirely distinct functional roles in sequence-dependent endosomal cargo sorting (Steinberg et al., 2013; McNally et al., 2017), they share structurally similar PX and FERM domains (Ghai et al., 2013; Ghai et al., 2015) (Figure 4A). A comparative immuno-isolation of GFP-SNX17 and GFP-SNX27 from HEK-293T cells established that SNX17 does not bind to ESCPE-1 (Figure 4B), consistent with the loss of ESCPE-1 binding to the SNX27 chimera containing the SNX17 FERM domain (Figure 2C). To identify potential regions for aDLF motif binding to the SNX27 FERM domain, we initially performed a comparative structural analysis between the X-ray structure of the SNX17 FERM domain (Ghai et al., 2013) and a homology model of the SNX27 FERM domain (Ghai et al., 2014). In particular we focused on the distribution and clustering of complementary basic amino acids. This revealed that the FERM domain of SNX27 possesses a basic surface defined in part by R437, R438, K495, K496, R498, and K501, residues that in SNX17 often corresponded to uncharged residues or negatively charged amino acids (Figure 4A and 4C). Indeed, charge swap site-directed mutagenesis of these residues generated full-length GFP-SNX27 mutants that when expressed and immuno-isolated from HEK-293T cells lacked the ability to associate with ESCPE-1; all mutants however retained the ability to associate with Retromer (Figure 4D). In direct ITC-quantified binding assays human SNX27(K495D), SNX27(R498D), and SNX27(K501D) all displayed no detectable binding to the human SNX1 75 to 92 peptide, while the SNX27(R496D) mutant displayed a reduce binding affinity of 56.4 µM when compared with the 12.7 µM observed for wild-type SNX27 (Figure 4E).

**Figure 4.**
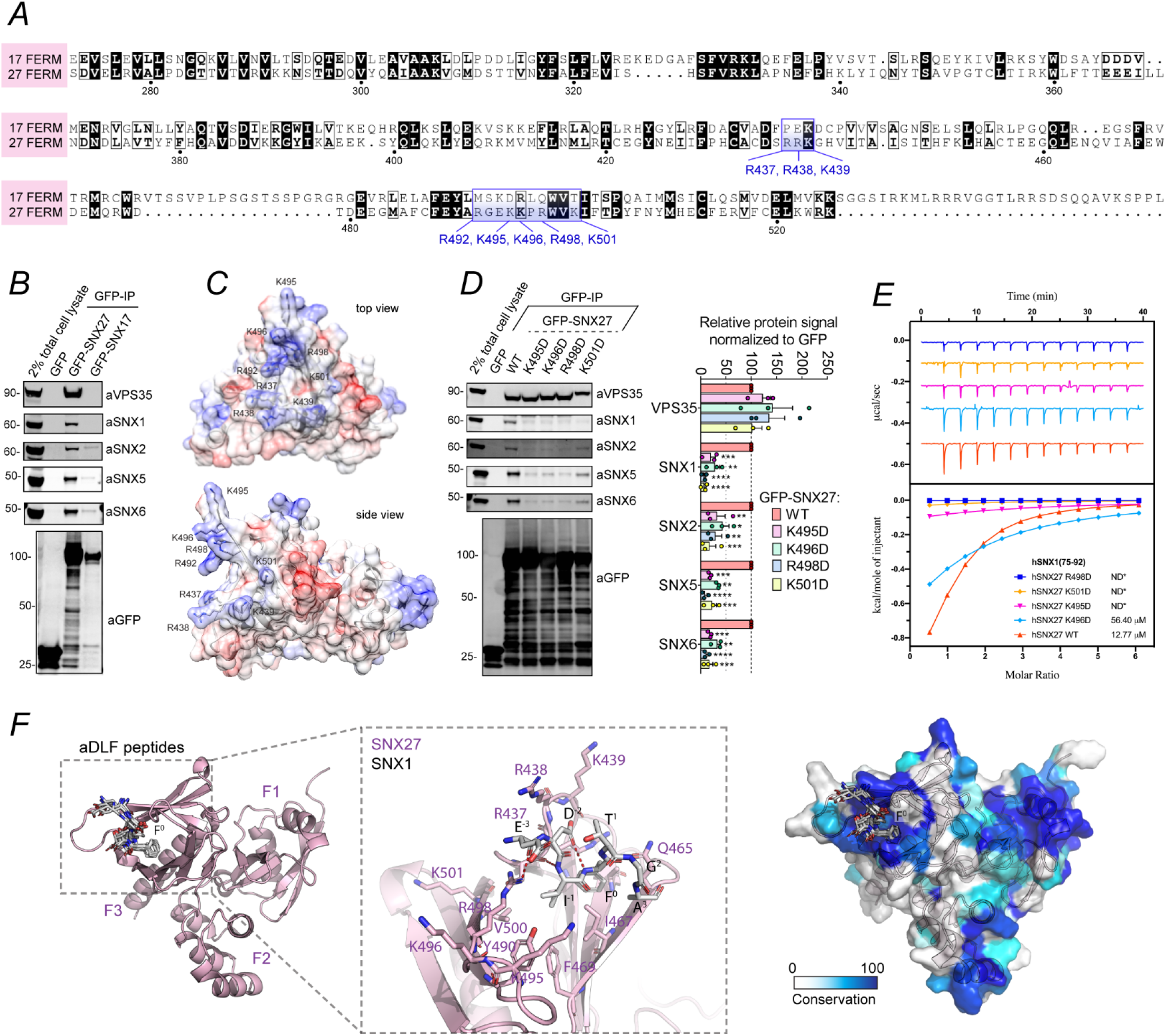
SNX1 and SNX2 N-termini bind a unique basic surface of the SNX27-FERM domain. **A** Sequence alignment of SNX27 FERM domain with the SNX17 FERM domain. The positively charged residues that constitute a basic surface unique to SNX27 are boxed and labelled in blue. **B** Co-immunoprecipitation of GFP-tagged SNX27 and SNX17. The constructs were transiently transfected in HEK-293T cells, the cell lysates were subjected to GFP-trap based immunoprecipitation and the immunoprecipitates were blotted for VPS35, SNX1, SNX2, SNX5, SNX6 and GFP. The blot is representative of three independent GFP traps. **C** Homology model of the SNX27 FERM domain (Ghai et al., 2015). The residues that are unique to SNX27 FERM domain and those that may account for the interaction with ESCPE-1 are labelled. **D** Co-immunoprecipitation of GFP-tagged SNX27 wild type (WT) and a panel of amino acid swap mutants across the FERM domain. The constructs were transiently transfected in HEK293T cells, the cell lysates were subjected to GFP-trap based immunoprecipitation and the immunoprecipitates were blotted for VPS35, SNX1, SNX2, SNX5 and SNX6. The band intensities were measured from n= 3 independent experiments using Odyssey software. The band intensity, normalized to GFP expression, is presented as the average fraction of the WT. The binding of SNX27 point mutants to VPS35, SNX1, SNX2, SNX5 and SNX6 was compared with the SNX27 WT using a two-way ANOVA and Dunnett’s test. **E** Binding of SNX1_75-92_ peptide to hSNX27_FL_ proteins by ITC (hSNX27 R437D: green; hSNX27 R498D: dark-blue; hSNX27 K501D: orange; hSNX27 K495D: lilac; hSNX27 K496D: sky-blue; hSNX27 WT: red-orange; ND*: no binding detected). The ITC graphs represent the integrated and normalized data fit with 1:1 ratio binding. The binding affinity (*K*_d_) is given as mean of at least two independent experiments. Thermodynamic parameters for the binding are provided in Supplementary Table 1. **F** The left panel shows the AlphaFold2 generated model of the SNX27 FERM domain (pink ribbons) bound to the core aDLF sequences (white sticks) of SNX1 and SNX2 with identical binding orientations. For clarity only one SNX27 model is shown with the eight aDLF peptides overlaid (two peptide models each of the four sequences from SNX1 and SNX2). The three subdomains of the SNX27 FERM domain F1, F2 and F3 are indicated. The middle panel shows a close-up view of the modelled SNX1 aDLF core sequence ^44^EDIFTGA^50^. The strictly conserved Phe sidechain of the aDLF sequences is numbered F^0^ for reference, and other residues are numbered with respect to this. The right panel shows the SNX27 surface coloured according to sequence conservation. The peptides are predicted to bind to a highly conserved pocket in SNX27. The ITC graphs represent the integrated and normalized data fit with 1:1 ratio binding. The binding affinity (*K*_d_) is given as mean of at least two independent experiments. Thermodynamic parameters for the binding are provided in Supplementary Table 1. Molecular masses are given in kilodaltons. Bars, error bars and symbols represent the mean, s.e.m. and individual data points, respectively. *P< 0.05, **P< 0.01, ***P< 0.001, ****P< 0.0001

Finally, we modelled the binding of the aDLF sequences from SNX1 and SNX2 with SNX27 using the AlphaFold2 machine learning algorithm (Jumper et al., 2021; Mirdita et al., 2021). In total we generated twelve models, with three models each of SNX27 in the presence of the four separate aDLF peptides (Figure 4F and Supplementary Figure 2). Without any prior inputs apart from the SNX27 and peptide sequences AlphaFold2 produced very high confidence models of the SNX27 FERM domain. Remarkably in all twelve models the aDLF peptides associated specifically with the SNX27 F3 subdomain, near the basic surface identified by mutagenesis (Figure 4F). Closer inspection showed that in eight of the models the core D[L/I]F residues adopted essentially identical bound structures, with the hydrophobic [L/I]F sidechains inserting into a hydrophobic pocket and interacting with SNX27 Ile-467, Phe-469, Tyr-490, and Val-500 and the acidic Asp sidechain surrounded by the complementary basic SNX27 residues Arg-437, Arg-438, Lys-439 and Arg-498 (Figure 4F). These models are entirely consistent and strongly support the mutagenesis data for the SNX27-peptide interactions.

### SNX27 association to ESCPE-1 couples cargo retrieval with promotion of cell surface recycling

Previously we established that by binding to the PDZ-binding motif present within the cytoplasmic facing tail of GLUT1, the SNX27-Retromer complex serves to retrieve internalized GLUT1 away from lysosomal mediated degradation (Steinberg et al., 2013, and see Figure 1A). To define the functional significance of the association of SNX27 to ESCPE-1 we therefore performed a series of rescue experiments to quantify the SNX27-dependent retrieval of internalized GLUT1 from lysosomal mediated degradation (Figure 5A). In the SNX27 knock out HeLa cells, GLUT1 displayed an enhanced rate of lysosomal mediated degradation following 8 hours of treatment with cycloheximide, data that are consistent with the loss of SNX27 expression leading to a reduced ability for transporter retrieval from the lysosomal degradative fate (Figure 5A). This phenotype was rescued by re-expression of wild-type GFP-tagged SNX27 (Figure 5A). Interestingly, expression of GFP-SNX27(R498D), a mutant that fails to associate with ESCPE-1 but retains binding to Retromer (Figure 4D and 4E), also fully rescued the retrieval of GLUT1 from lysosomal degradation (Figure 5A). The coupling of SNX27-Retromer to the ESCPE-1 complex is therefore not required for the retrieval of GLUT1 from lysosomal degradation.

**Figure 5.**
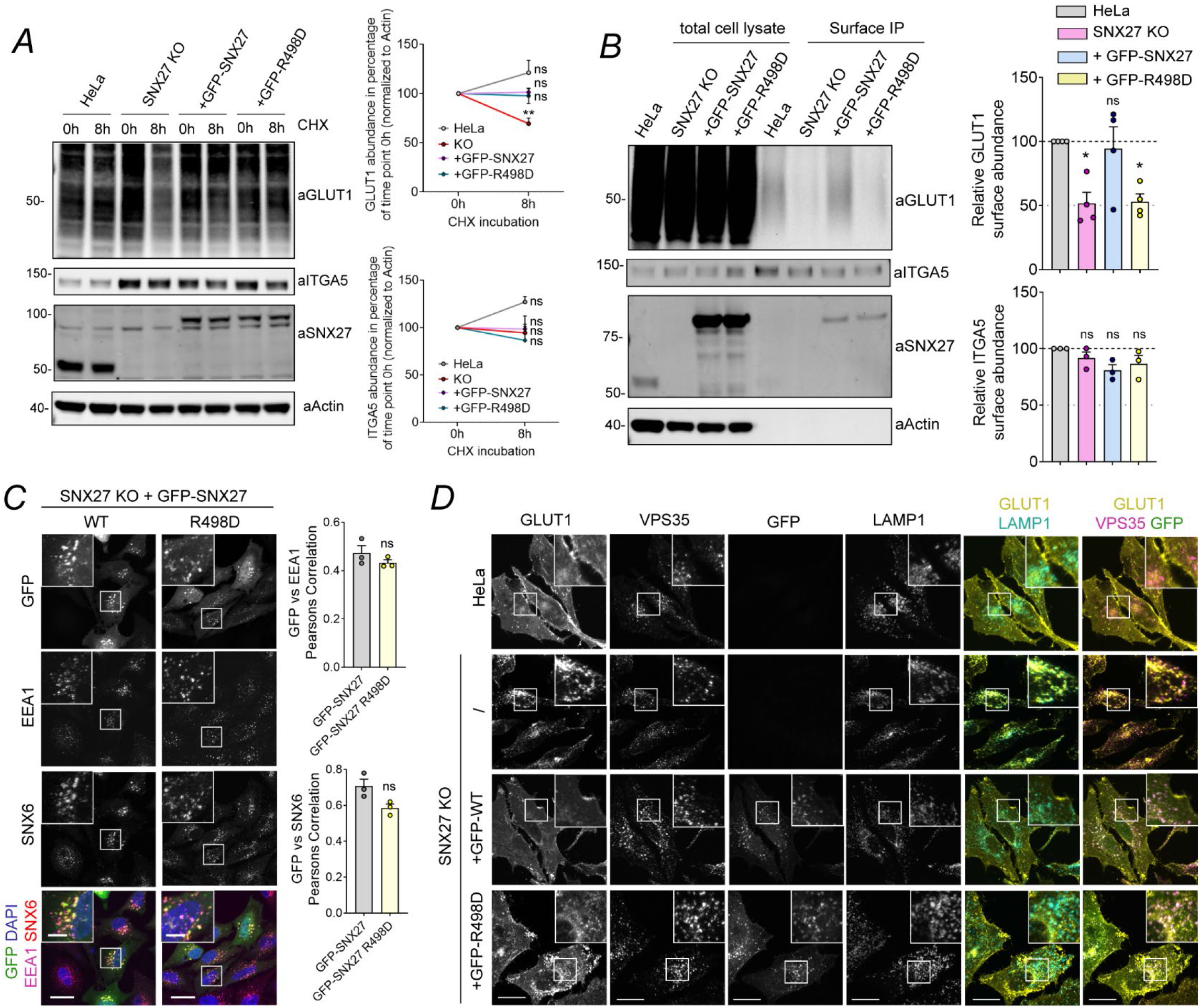
SNX27:ESCPE-1 interaction is required for endosome-to-plasma membrane recycling of SNX27 cargos. **A** Degradation assay in parental HeLa cells or SNX27 KO HeLa cells that were transiently transfected with GFP-SNX27 or GFP-SNX27(R498D) mutant. 48h after transfection, cells were incubated with 10 µg/ml cycloheximide and lysed at different time points as indicated. The band intensities of endogenous GLUT1 and ITGA5 were measured from n= 3 independent experiments using Odyssey software. The levels of GLUT1 and ITGA5 at 8h were compared with the corresponding level at 0h timepoint. Analysis was done using a two-way ANOVA and Sidak’s test. GLUT1: P = 0.0735 (HeLa), 0.0068 (KO), 0.9997 (+GFP-SNX27), 0.9975 (+GFP-R498D). ITGA5: P = 0.1151 (HeLa), 0.9820 (KO), 0.9999 (+GFP-SNX27), 0.6899 (+GFP-R498D). **B** Analysis of surface levels of GLUT1 and ITGA5 in parental HeLa cells or SNX27 KO HeLa cells that were transiently transfected with GFP-SNX27 or GFP-SNX27(R498D) mutant. 48h after transfection, cells were subjected to surface biotinylation followed by streptavidin-based immunoisolation. The immunoisolates where blotted for GLUT1 and the band intensities were measured from n = 4 independent experiments using Odyssey software and compared to the levels in parental HeLa using one-way ANOVA and Dunnett’s test. GLUT1: P = 0.0134 (KO vs HeLa), 0.9587 (+GFP-SNX27 vs HeLa), 0.0163 (+GFP-SNX27 R498D vs HeLa). The immunoisolates where blotted for ITGA5 and the band intensities were measured from n = 3 independent experiments using Odyssey software and compared to the levels in parental HeLa using one-way ANOVA and Dunnett’s test. ITGA5: P = 0.5616 (KO vs HeLa), 0.0733 (+GFP-SNX27 vs HeLa), 0.2354 (+GFP-SNX27 R498D vs HeLa). **C** Representative images of SNX27 KO HeLa cells transiently transfected with GFP-SNX27 or GFP-SNX27(R498D). 48h after transfection, cells were fixed and immunostained for the endosomal markers EEA1 and SNX6. Magnified views of the white boxes are shown boxed. Cell numbers analysed for colocalization were: 60 GFP–SNX27 and 60 GFP–SNX27 R498D cells across n = 3 independent experiments. The Pearson’s coefficient values were compared using a two-tailed unpaired t-test; for EEA1 P = 0.2971, SNX6 P = 0.0506. **D** Representative images of parental Hela cells, SNX27 KO HeLa cells transiently transfected with GFP-SNX27 or GFP-SNX27(R498D). 48h after transfection, cells were fixed and immunostained for the cargo GLUT1, the endosomal marker VPS35, and the late endosome/lysosome marker LAMP1. Scale bars, 25 μm (micrographs) and 5 μm (magnified images). Molecular masses are given in kilodaltons. Bars, error bars and symbols represent the mean, s.e.m. and individual data points, respectively. *P< 0.05, **P< 0.01, ***P< 0.001, ****P< 0.0001, ns = not significant.

Next, we turned to examining the promotion of GLUT1 recycling back to the cell surface. Here we utilized restricted cell surface biotinylation to quantify the amount of GLUT1 at the cell surface (Steinberg et al., 2013). Compared with control HeLa cells, SNX27 knock out HeLa cells displayed a reduced cell surface abundance of GLUT1 as a result of missorting of internalized GLUT1 for lysosomal degradation (Figure 5B). Again, this phenotype was rescued by expression of GFP-SNX27 (Figure 5B) (Steinberg et al., 2013). In contrast, expression of GFP-SNX27(R498D) failed to rescue the cell surface level of GLUT1 (Figure 5B), consistent with the association of SNX27 with ESCPE-1 being required to promote the cell surface recycling of internalized GLUT1. Importantly, GFP-SNX27(R498D) exhibited an endosomal localization indistinguishable from wild-type SNX27, consistent with the inability to promote endosome-to-plasma membrane recycling of GLUT1 not arising from a loss of endosomal localisation (Figure 5C). Furthermore, confocal analysis of the steady-state distribution of GLUT1 in GFP-SNX27(R498D) rescue cells confirmed that GLUT1 was retrieved from the lysosomal degradative fate, as shown by reduced co-localisation with the LAMP1-labelled lysosome (Figure 5D). However, unlike the rescue with wild-type GFP-SNX27, where GLUT1 was enriched at the cell surface, the GLUT1 in the GFP-SNX27(R498D) rescue became ‘trapped’ in a Retromer labelled endosomal compartment and failed to repopulate the cell surface (Figure 5D). Together, these data are consistent with SNX27 orchestrating cargo retrieval from the degradative fate through its PDZ domain-mediated association with cargo and Retromer, while the SNX27 FERM domain mediates association with ESCPE-1 to promote tubular-based plasma membrane recycling of the retrieved cargo (Figure 6A).

**Figure 6.**
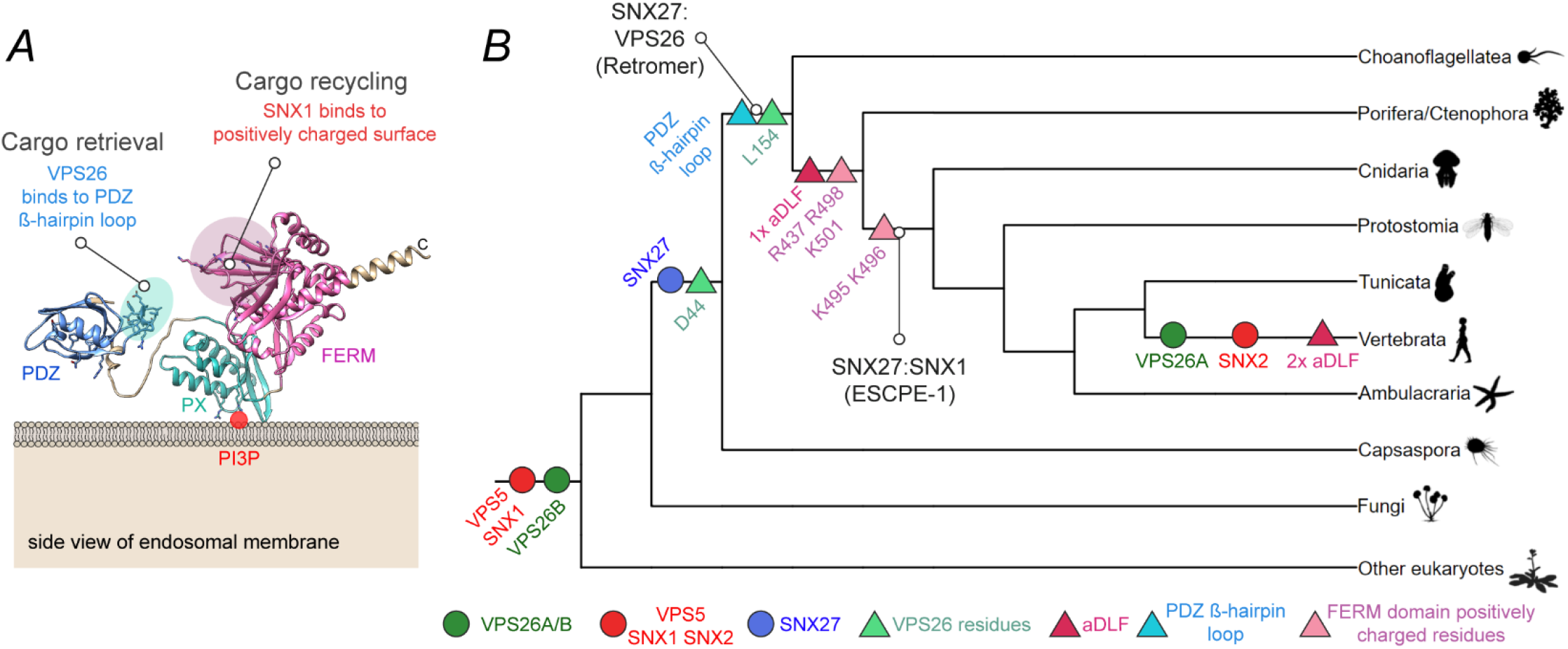
Molecular evolution of the SNX27-Retromer and SNX27-ESCPE-1 interactions. **A** Alphafold2 structural prediction for full length human SNX27 (Tunyasuvunakool et al., 2021). The residues of the exposed ß-hairpin loop in the SNX27 PDZ domain (67-79) that account for the interaction with Retromer are highlighted in cyan. Those residues of the FERM domain that account for the interaction with ESCPE-1 (positively charged surface) are highlighted in pink. Our data support a model where the interaction between SNX27 and Retromer is required for the retrieval of PDZ cargos away from the degradative fate. Subsequently, the interaction between SNX27 and ESCPE-1 is required for the export and endosome-to-plasma membrane recycling of PDZ cargos. **B** A cartoon representation incorporating the results of multiple phylogenetic analyses, including VPS5/SNX1/SNX2, SNX27 and VPS26A/VPS26B (see Supplementary Figure 3A for detailed phylogenies). The gene families that contain VPS5/SNX1/SNX2 and VPS26A/VPS26B are present across eukaryotes: SNX1 and SNX2, and VPS26A and VPS26B, arose by gene duplication in the common ancestor of vertebrates. SNX27 is found in metazoans, choanoflagellates and the filasterean *Capsaspora*, but not in more distantly related opisthokonts such as Ichthyosporea or Fungi; it is therefore likely arose in the filazoan common ancestor. The residue D44 (VPS26B, *Homo sapiens*) in VPS26 is found across Filazoa, laying the foundations for the binding interaction between VPS26 and SNX27. L154 (VPS26B, *Homo sapiens*) likely arose in the ancestral choanozoan along with the PDZ loop and key interacting residues (67-97 L67, L74, SNX27, *Homo sapiens*) suggesting the SNX27-Retromer interaction was already present in the common ancestor of metazoans and choanoflagellates. Key residues of the FERM domain: R437, R498, K501 (SNX27, *Homo sapiens*) are recognizable in the ancestral metazoan. The first aDLF motif (SNX1) was acquired in the common ancestor of cnidarians and bilaterians, along with the residues K495 and K496 which make up part of the FERM domain in SNX27, potentially suggesting that SNX27:ESCPE-1 binding has evolved by this time, if not earlier during metazoan evolution. Interestingly, the first aDLF motif does not appear to be present in the placazoan or ctenophoran sequences, (although the R437, R498 and K501 residues are present), which might reflect secondary loss of the motif in these groups. The second aDLF motif is only present in vertebrates.

### Evolutionary assembly of the SNX27:Retromer:ESCPE-1 sorting axis

We used phylogenetic analyses to investigate the evolution of the association of SNX27 with Retromer and with the ESCPE-1 complex. The inferred phylogeny of SNX27 suggests the gene was ancestrally present in filozoans (that is, the group comprising filastereans, choanoflagellates and metazoans) (Supplementary Figure 3A). In SNX27 an exposed ß-hairpin in the SNX27 PDZ domain, encoded by residues 67-79 in human SNX27, engages a groove in the VPS26A (and VPS26B) subunit of Retromer (Gallon et al., 2014) (Figure 6A). An equivalent sequence, including key residues Leu-67 and Leu-74 of the ß-hairpin, is present in SNX27 across all choanozoans (this is, the group comprising metazoans and choanoflagellates; Figure 6B and Supplementary Figure 3A). Key residues in VPS26A required for SNX27 binding include Asp-44 and Leu-154, equivalent residues in VPS26B being Asp-42 and Leu-152 (Gallon et al., 2014). The inferred phylogeny of VPS26A and VPS26B, revealed that VPS26B is ancestrally present across eukaryotes, with a duplication of VPS26B in vertebrates leading to the emergence of VPS26A (Figure 6B, Supplementary Figure 3B). Interestingly, within VPS26B the Asp-44 residue is present ancestrally in filozoans, while the Leu-154 residue evolved in choanozoans (Figure 6B). Together these analyses suggest that the SNX27-Retromer association first appeared in the last common ancestor of choanoflagellates and metazoans.

The association between SNX27 and ESCPE-1 depends on the presence of binding motifs in SNX1 and SXN2 and the FERM domain of SNX27. SNX1 is present across eukaryotes; a gene duplication giving rise to SNX1 and SNX2 from a single ancestral protein occurred in the common ancestor of vertebrates (Figure 6B, Supplementary Figure 3C). Several motifs required in the FERM domain of *Homo sapiens* SNX27 for binding to ESCPE-1 appear to have evolved in the common ancestor of metazoans or early during their radiation (Figure 6B); many of the key residues (Arg-437, Arg-498, Lys-501) were in place in the last metazoan common ancestor or prior to the divergence of cnidarians and bilaterians (Lys-495,Lys-496). For SNX1, the first aDLF motif was likely already present in the last metazoan common ancestor: it is present in sequences from bilaterians and Porifera (sponges) but is absent in cnidarians, placazoans and ctenophorans, which might represent secondary losses of the motif. As for the second aDLF motif, this is present in all vertebrates. This analysis therefore suggests that the SNX27:ESCPE-1 interaction evolved after the SNX27-Retromer link during early metazoan evolution.

## DISCUSSION

Here in combining biochemical, structural and cellular analyses we have established that acidic-Asp-Leu-Phe (aDLF) motifs in the redundant SNX1 and SNX2 subunits of ESCPE-1 directly associate with the FERM domain of SNX27 within the SNX27-Retromer assembly. Studies of endosomal cargo sorting show that this association is necessary to promote the ESCPE-1-dependent tubulovesicular cell surface recycling of internalised transmembrane proteins. Our data suggests that ESCPE-1 mediated recycling occurs subsequently or in conjunction with SNX27-Retromer-dependent retrieval from lysosomal degradation. Building on previous studies (Temkin et al., 2011; Steinberg et al., 2013; Gallon et al., 2014), these data collectively identify the SNX27:Retromer:ESCPE-1 assembly as a dynamic endosomal coat complex that couples sequence-dependent cargo recognition with membrane remodeling to facilitate the formation of cargo-enriched transport carriers for cargo recycling. In addition, we have begun to describe the functional diversification of Retromer as it has evolved from its tight association with the SNX-BAR homologues Vps5 and Vps17, as originally identified in yeast (Seaman, et al., 1998), to a more transient engagement of ESCPE-1 in metazoans and their relatives mediated at least in part through the SNX27 cargo adaptor.

At present we propose the following model to describe the functional connectivity of the SNX27:Retromer:ESCPE-1 coat complex within the context of cargo retrieval and recycling. Newly endocytosed cargo containing a functional SNX27 PDZbm are recognized by SNX27, associated with the PI(3)P-enriched early endosome, and these interaction together can stabilize the endosomal residency of SNX27 (Ghai et al., 2013). As the endosome matures, SNX27 binds to Retromer through direct association of VPS26 to the SNX27 PDZ domain. These interactions are mutually stabilizing, thereby locking down cargo association (Gallon et al., 2014; Clairfeuille et al., 2016): this assembly is essential for cargo retrieval from the lysosomal degradative fate (Steinberg et al., 2013). Retromer dimerization and the formation of Retromer arches (Kovtun et al., 2018; Kendall et al., 2020; Leneva et al., 2021) may aid SNX27 and cargo clustering thereby forming the retrieval sub-domain and initiating weak membrane curvature (Simunovic et al., 2015). These events are supported through the Retromer-mediated recruitment of the WASH complex and the localized production of branched filamentous actin (Harbour et al., 2012; Jia et al., 2012). An additional accessory protein, ANKRD50, may serve to stabilize the formation of the SNX27:Retromer:WASH assembly (Kvainickas et al., 2017). To this emerging ‘bud’ ESCPE-1 is recruited through recognition of 3-phosphoinositides and membrane curvature (Cozier et al., 2002; Carlton et al., 2004; Chandra et al., 2019), and the direct binding to SNX27 described in the present study. The timed increase in ESCPE-1 residency and localized concentration provides the hand-over mechanism to ensure that the captured and enriched cargo enter the forming tubular profile and the ensuing transport carrier. Continued localized WASH-mediated actin polymerization and the recruitment of motor proteins facilitates tubule maturation and fission (Wassmer et al., 2009; Hong et al., 2009; Freeman et al., 2014) (Figure 7). Additional cargo entry may be achieved through direct recognition by ESCPE-1 and yet to be described mechanisms (Simonetti et al., 2019; Yong et al., 2020). As with the assembly of other coat complexes (Schmid and McMahon, 2007), association of the SNX27:Retromer:ESCPE-1 coat is driven by avidity through a series of low affinity interactions that evoke the consideration of checkpoints for monitoring the fidelity of pathway progression. It is important to state that this model is certainly an over-simplification and many key questions remain unanswered. Not least whether such a coat complex can be visualized through membrane associated cryo-electron tomography of reconstituted SNX27:Retromer:ESCPE-1 perhaps in the presence of ANKRD50.

**Figure 7.**
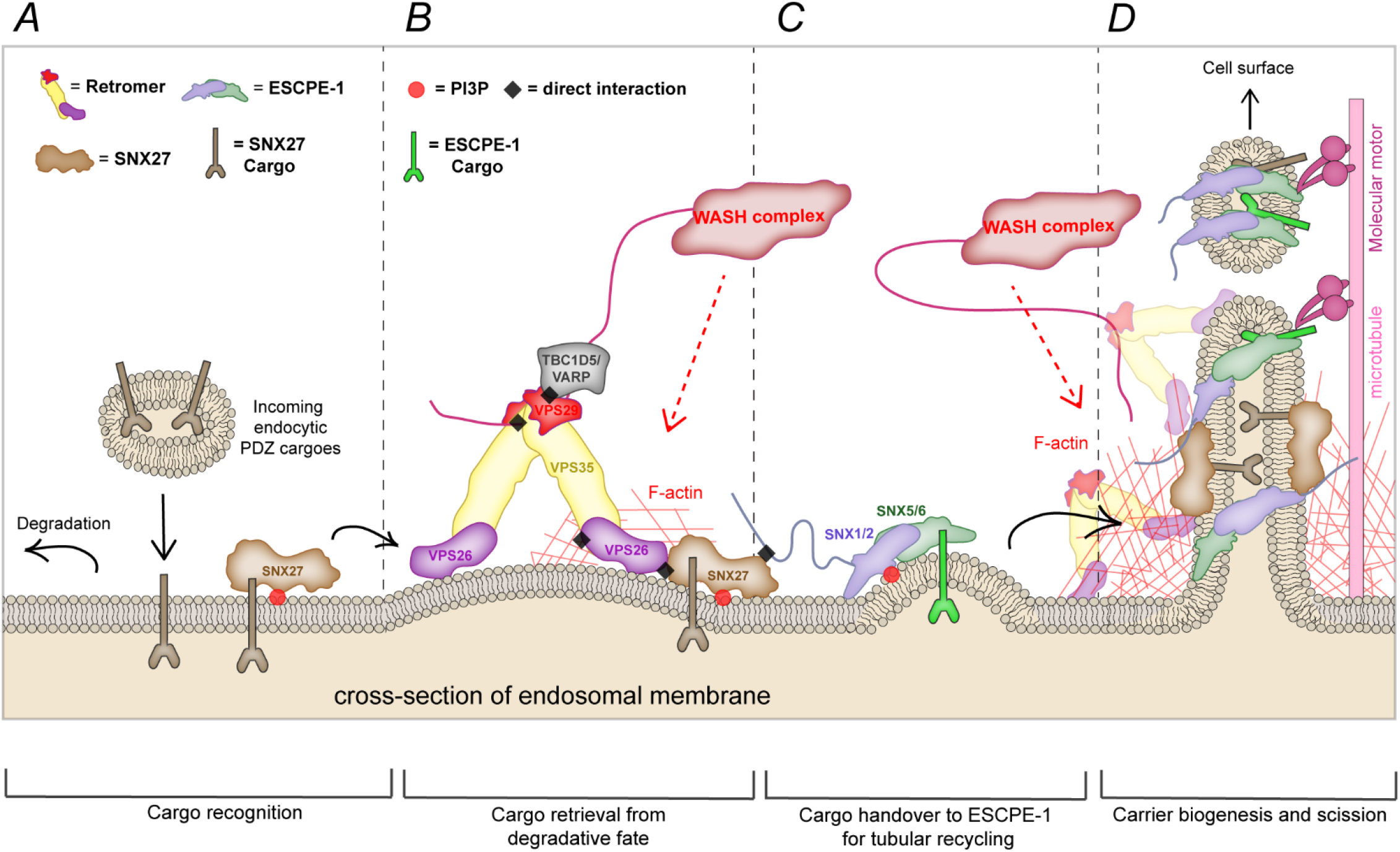
A two-step model for the endosomal sorting of SNX27 cargos. **A** Schematics of the endosomal sorting of PDZ cargos. Cargos possessing a PDZbm that enter the endosomal system are recognized by SNX27 via its PDZ domain. **B** SNX27 binding to Retromer aids the retrieval of cargos from the lysosomal degradative path. **C** Once cargos have been retrieved, the SNX27 FERM engages the N-terminus of SNX1 and SNX2 mediating the inclusion of SNX27 cargos into ESCPE-1 tubulovesicular carriers. **D** This mechanism results in SNX27 cargos entering carriers that exit endosomes and are transported, directly and/or indirectly, to the cell surface.

The importance of the N-terminal extension of SNX1 and SNX2 have not previously been considered. While we have established that they constitute primary sites of interaction with SNX27, it is tempting to speculate that they could orchestrate a broader set of protein-protein interactions. Interestingly, these unstructured N-termini may extend to over 30 nm in length and could allow the ESCPE-1 oligomeric assemblies to capture peripheral factors for the fine-tuned regulation of tubular maturation and scission. This would resemble the organization of the endocytic clathrin coat where several accessory proteins protrude from the core clathrin lattice into the cytoplasm to enable a regulated function of the endocytic coat during endocytic vesicle formation (Skruzny et al., 2020).

Other than mediating the recruitment of accessory proteins, there is some evidence that these N-terminal extensions could play a role in the assembly and membrane targeting of SNX-BAR dimers. In fact, besides SNX1 and SNX2, SNX4, SNX7, SNX8 and SNX30 all possess unstructured N-terminal extensions of different length and of unknown function. The low-complexity region of the SNX8 homologue Mvp1 appears to stabilize soluble Mvp1 in a cytosolic tetrameric form where the PX and BAR domains are masked and unable to target membranes (Sun et al., 2020). The release of the N-terminus liberates functional dimers of Mvp1 facilitating their membrane association and function (Sun et al., 2020). It will be interesting to explore whether a similar mechanism also regulates ESCPE-1 targeting to the endosomal network in mammalian cells during endosomal recycling.

An important aspect of cargo sorting by SNX27 is regulation by post-translational modification. Phosphorylation of residues within the core C-terminal PDZbm negatively impairs SNX27 association, but phosphorylation of residues preceding the minimal PDZbm can dramatically enhance affinity for the SNX27 PDZ domain (Clairfeuille et al., 2016). A well characterized example of cargo tail phosphorylation modulating SNX27 affinity and subsequent endosomal sorting is the β2AR (Cao et al., 1999; Clairfeuille et al., 2016). Furthermore, phosphorylation of SNX27 at Ser-51 alters the conformation of its PDZbm-binding pocket decreasing the affinity for cargo proteins, thereby inhibiting endosomal sorting (Mao et al., 2021) and Ser-302 and Ser-304 phosphorylation in VPS26B can modulate SNX27 association (Wang et al., 2021). Similarly, ESCPE-1 function can be regulated by phosphorylation, in this case Ser-226 of SNX5 undergoes phosphorylation which decreases dimerization with SNX1 and SNX2 and reduces endosomal recycling (Itai et al., 2018). Therefore, a fascinating question regards whether the SNX27:ESCPE-1 interaction may be orchestrated by post-translational modification. We have here shown that a series of acidic residues upstream of the DLF motif in the SNX1 and SNX2 N-termini contribute to the binding to SNX27 (Figure 3F). Interestingly, several phosphorylation sites in the aDLF sequences of SNX1 have been annotated, including Ser-39, Ser-41 and Ser-72, as well as Thr-72 (Dephoure et al., 2008; Zhou et al., 2013; Bian et al., 2014). It is tempting to speculate that multiple phosphorylations of the ESCPE-1 N-termini may regulate its association with SNX27-Retromer, and in turn dictate the timing and kinetics of the endosomal recycling route.

Finally, a fascinating question relates to the molecular evolution of Retromer (Koummandou et al., 2011). In yeast, Retromer can form a stable pentameric assembly of two sub-complexes, the so called ‘cargo-selective’ Vps26:Vps35p:Vps29p heterotrimer and the membrane remodeling Vps5p and Vps17p heterodimer (Seaman et al., 1998); the latter being considered analogous to ESCPE-1. Two functional processes, sequence-dependent cargo recognition and the biogenesis of tubulovesicular transport carriers, are therefore brought together in a single pentameric assembly (Suzuki et al., 2019). In metazoans the VPS26:VPS35:VPS29 Retromer and ESCPE-1 appear to no longer form a stable pentameric assembly (Norwood et al., 2011), rather they have diverged into two separate complexes and in so doing have expanded their functional roles (Simonetti et al., 2017; Kvanicikas et al., 2017; Simonetti et al., 2019; Yong et al., 2020). Our phylogenetic analysis suggests that the SNX27:Retromer:ESCPE-1 assembly observed in metazoans evolved in a stepwise manner, with the SNX27-Retromer interaction evolving prior to the origin of metazoans, in the common ancestor of metazoans and choanoflagellates – the group of unicellular and colonical flagellates that are the closest living relatives of animals. The SNX27:ESCPE-1 interaction evolved later, during early metazoan evolution and likely prior to the divergence of cnidarians and bilaterians. Thus, within the SNX27:Retromer:ESCPE-1 pathway the retrieval of cargo from the degradative fate appears to have evolved prior to the direct coupling to ESCPE-1 for promotion of tubulovesicular-based recycling to the cell surface. The divergence of the ancestral pentameric Retromer coat complex therefore appears to have occurred to accommodate the expanding complexity of the endomembrane system in metazoans (Rout and Field, 2017). The ability of metazoan Retromer to associate with SNX27 and the ability of ESCPE-1 to directly bind cargo through a helix-loop-helix region (Simonetti et al., 2019; Yong et al., 2020), features not observed in yeast, have greatly expanded the capacity for sequence-dependent endosomal retrieval and recycling in metazoans. Indeed, it is tempting to speculate that the increased plasticity of the SNX27:Retromer:ESCPE-1 axis, and the existence of additional Retromer-independent pathways (Cullen and Steinberg, 2018), has allowed the metazoan endosomal network to accommodate the evolving diversity and number of cargos undergoing sequence-dependent sorting, leading to an expansion in the number of recycling itineraries that cargos can engage to interface with distinct membrane compartments. Dissecting how these sorting complexes and pathways are integrated and regulated, for example by post-translational modifications, in order to shape sequence-dependent cargo sorting within the metazoan endosomal network will be a major challenge moving forward.

Overall, in defining the mechanistic basis of SNX27-Retromer association with ESCPE-1 we have provided new functional insight into the endosomal sorting of hundreds of internalized PDZbm-containing receptors, channels, transporter, enzymes and adhesion molecules for repopulation of the cell surface (Steinberg et al., 2013). This increased molecular understanding provides new insight into the role of sequence-dependent endosome sorting in cellular processes ranging from the establishment of polarity and the uptake of nutrients through to synaptic function and cancer cell metastasis in both health and disease (Chandra et al., 2021).

## MATERIAL AND METHODS

### Antibodies

Antibodies used in this study were: mouse monoclonal antibodies GFP (clones 7.1 and 13.1; 11814460001; Roche) (1:2000 for WB, 1:400 for IF), SNX1 (clone 51/SNX1; 611482; BD) (1:1000 for WB, 1:200 for IF), SNX2 (clone 13/SNX2; 5345661; BD) (1:1000 for WB, 1:200 for IF), SNX6 (clone d-5,365965; Santa Cruz Biotechnology, Inc.) (1:1000 for WB, 1:200 for IF), SNX27 (ab77799, Abcam) (1:1000 for WB, 1:200 for IF), β-actin (A1978; Sigma-Aldrich) (1:2000 for WB), ITGA5 (610633, BD) (1:1000 for WB); rabbit monoclonal antibodies: VPS35 (EPR11501(B); 157220; Abcam) (1:1000 for WB), GLUT1 (ab115730, Abcam) (1:1000 for WB, 1:200 for IF); rabbit polyclonal antibodies: VPS26A (23892; Abcam) (1:1000 for WB), VPS35 (97545; Abcam) (1:1000 for WB, 1:200 for IF), VPS29 (98929; Abcam) (1:100 for WB); goat polyclonal antibodies to VPS35 (10099; Abcam) (1:200 for IF), GFP rabbit polyclonal (GTX20290; GeneTex) (WB 1:2000), EEA1 rabbit monoclonal (C45B10; Cell Signaling Technologies) (IF 1:200), LAMP1 mouse monoclonal (clone H4A3, AB2296838; DSHB) (IF 1:200).

### Cell culture and transfection

HeLa and HEK-293T cell lines were sourced from ATCC. Authentication was from the ATCC. Cells were grown in DMEM (Sigma-Aldrich) supplemented with 10% (v/v) FCS (Sigma-Aldrich) and penicillin/streptomycin (Gibco) and grown under standard conditions. FuGENE HD (Promega) was used for transient transfection of DNA according to the manufacturer’s instructions. Cycloheximide was used to prevent up-regulation of protein synthesis. In this case, cycloheximide (C7698; Sigma-Aldrich) at 10 µg/ml was added to the cells 48 h after FuGENE transfection for the indicated time points. For the generation of CRISPR-Cas9 SNX27 knock-out HeLa cells, HeLa cells were transfected with pX330 plasmid coding for the gRNA against the gene of interest together with a puromycin resistance–expressing plasmid. Cells were then subjected to puromycin selection for 24 h. subjected to lysis and Western blotting to determine the levels of the target proteins. The VPS35 knock-out clonal cell line used in this study was characterized previously (Kvainickas et al., 2017). For GFP-based immunoprecipitations, HEK293T cells were transfected with GFP constructs using polyethylenimine (Sigma-Aldrich) and expression was allowed for 48 h.

### Immunoprecipitation and quantitative Western blot analysis

For western blotting, cells were lysed in PBS with 1% (v/v) Triton X-100 and protease inhibitor cocktail. The protein concentration was determined with a BCA assay kit (Thermo Fisher Scientific), and equal amounts were resolved on NuPAGE 4–12% precast gels (Invitrogen). Blotting was performed onto polyvinylidene fluoride membranes (Immobilon-FL; EMD Millipore) followed by detection using the Odyssey infrared scanning system (LI-COR Biosciences). For GFP-based immunoprecipitations, HEK-293T cells were lysed 48 h after transfection in immunoprecipitation buffer (50 mM Tris-HCl, 0.5% (v/v) NP-40, and Roche protease inhibitor cocktail) and subjected to GFP trap (ChromoTek). Immunoblotting was performed using standard procedures. Detection was performed on an Odyssey infrared scanning system (LI-COR Biosciences) using fluorescently labeled secondary antibodies. In using the Odyssey, we routinely performed Western blot analysis where a single blot is simultaneously probed with antibodies against two proteins of interest (distinct antibody species) followed by visualization with the corresponding secondary antibodies conjugated to distinct spectral dyes.

### Biotinylation of cell surface proteins

For surface biotinylation experiments, fresh Sulfo-NHS-SS Biotin (Thermo Scientifics, #21217) was dissolved in ice-cold PBS at pH 7.8 at a final concentration of 0.2 mg / ml. Cells were washed twice in ice-cold PBS and placed on ice to slow down the endocytic pathway. Next, cells were incubated with the biotinylation reagent for 30 minutes at 4°C followed by incubation in TBS for 10 minutes to quench the unbound biotin. The cells were then lysed in lysis buffer and subjected to Streptavidin beads-based affinity isolation (GE-Healthcare).

### Immunofluorescence staining

Cells were fixed in 4% (v/v) PFA for 20 min and washed three times in PBS and permeabilized with 0.1% (v/v) Triton X-100. For LAMP1 staining, cells where instead permeabilized by submersion in liquid nitrogen for 20 seconds. Fixed cells were blocked in 1% (w/v) BSA and incubated in primary antibody and respective secondary antibody (Alexa Fluor; Thermo Fisher Scientific) in 1% (w/v) BSA.

### Image acquisition and image analysis

Microscopy images were collected with a confocal laser-scanning microscope (SP5 AOBS; Leica Microsystems) attached to an inverted epifluorescence microscope (DMI6000; Thermo Fisher Scientific). A 63× 1.4 NA oil immersion objective (Plan Apochromat BL; Leica Biosystems) and the standard SP5 system acquisition software and detector were used. Images were captured at room temperature as z stacks with photomultiplier tube detectors with a photocathode made of gallium-arsenide-phosphide (Leica Microsystems) for collecting light emission. Images were captured using Application Suite AF software (version 2.7.3.9723; Leica Microsystems) and then analyzed with the Volocity 6.3 software (PerkinElmer).

### Statistics and Reproducibility

All quantified Western blot are the mean of at least three independent experiments. Statistical analyses were performed using Prism 7 (GraphPad Software). Graphs represent means and S.E.M. For all statistical tests, P < 0.05 was considered significant and is indicated by asterisks.

### Molecular biology and cloning for recombinant protein production

For bacterial expression, full-length mouse SNX1 with an N-terminal His-tag (mSNX1_FL_ hereafter) was cloned into the pMW172Kan vector (Collins et al., 2008) and the truncated mSNX1 construct containing residues 1 to 139 was cloned into pGEX4T-2 vector for expression as a thrombin-cleavable N-terminal GST fusion protein. DNA encoding full-length mouse SNX27 (mSNX27_FL_ hereafter) was inserted into the pMCSG7 vector (Eschenfeldt et al., 2009) by the ligation independent cloning (LIC-cloning) approach. Human SNX27 constructs, hSNX27_FL_, hSNX27_FERM_, and hSNX27_FL_ single-site mutants R437D, K495D, K496D, R498D and K501D were cloned into the pET-28a vector by GenScript^®^ and codon optimised for bacterial protein expression. An additional TEV cleavage site was inserted between the N-terminal His-tag and hSNX27 coding sequences.

### Protein expression and purification

The DNA constructs of mSNX27 and mSNX1 were expressed using *Escherichia coli* BL21 CodonPlus^TM^ (DE3) competent cells, while the hSNX27 constructs were expressed using *E. coli* BL21 (DE3) competent cells. The cells were grown in Ultra Yield^TM^ flasks at 37 ℃, 180 rpm until the cell density reached OD_600_ ∼0.8. Protein expression was then induced with 1 mM isopropyl-β-D-1-thiogalactoside (IPTG) and cells incubated for 16 to 18 h at 18℃. For SNX27, 0.5 mM IPTG was added for induction to get optimal protein expression yield. To collect cell pellets, the cells were harvested by 10 min centrifugation (JLA 8.1 rotor) at 6000 rpm, 4℃.

For SNX1 purification, the cell pellet was resuspended in lysis buffer containing 50 mM HEPES pH 7.5, 200 mM NaCl, 5% glycerol, 1 mM β-mercaptoethanol (β-ME), 1 mM benzamidine and 10 μg/mL DNase I dissolved in phosphate-buffered saline (PBS) containing 10% glycerol. For SNX27 proteins, the cell pellet was resuspended in lysis buffer containing 20 mM Tris-HCl pH 8.0, 500 mM NaCl, 5% glycerol, 1 mM β-ME, 1% Triton^TM^ X-100, 1 mM benzamidine and 10 μg/mL DNase I dissolved in PBS containing 10% glycerol. Cells were lysed by high-pressure homogenisation (35 Kpsi). The homogenate was centrifuged for 30 min at 4℃ using a JLA 25.50 rotor with 19,000 rpm. The supernatant of His-tagged protein homogenates were loaded onto pre-equilibrated TALON^®^ metal affinity resin (Clonetech) or Ni-nitrilotriacetic acid (NTA) resin, while the supernatant of GST-fusion protein homogenates were loaded onto pre-equilibrated glutathione Sepharose^TM^ beads (GE healthcare), and incubated on an orbital rotator for an hour in cold room.

Affinity matrices with bound fusion proteins were loaded into empty glass chromatography columns (Bio-Rad) washed with 100 ml of washing buffer containing either 50 mM HEPES pH 7.5, 200 mM NaCl, 5% glycerol, 1 mM β-ME, or 20 mM Tris-HCl pH 8.0, 200 mM NaCl, 5% glycerol, 1 mM β-ME. For His-tagged protein purification, 10 mM and 300 mM imidazole was added to wash and elution buffers respectively. To collect the GST-tagged mSNX1 constructs, 50 mM reduced glutathione in elution buffer was applied for elution. cOmplete^TM^ EDTA-free protease inhibitor tablets were added to the eluted SNX1 samples to reduce proteolytic degradation of their disordered N-terminal region.

The fractions containing eluted proteins were concentrated to ∼5 ml by Amicon^TM^ ultrafiltration devices (10 kDa or 30 kDa molecular weight cut-off) (Millipore) and centrifuged in a bench centrifuge for 10 min at maximum speed to remove any precipitated material before being subjected to size-exclusion chromatography (SEC). For SEC, the HiLoad^®^ 16/600 Superdex 200 (GE Healthcare) column was used with proteins eluted in SEC buffer containing 50 mM HEPES pH 7.5, 200 mM NaCl, 5% glycerol, 1 mM β-ME or 20 mM Tris-HCl pH 8.0, 100 mM NaCl, 5% glycerol, 1 mM β-ME. To check the purify and integrity of the purified fractions, sodium dodecyl sulfate polyacrylamide gel electrophoresis (SDS-PAGE) was applied using Bolt^TM^ 12 % Bis-Tris gel with 1x MES running buffer at 190V for 25 min. After the SDS-PAGE gels were stained with Coomassie Brilliant Blue R250 solution for visualization. For hSNX27_FERM_, an additional anion exchange chromatography (IEX) was applied using the Mono Q™ anion exchange chromatography column pre-equilibrated with IEX buffer A: 20 mM Tris-HCl pH 8.0, 100 mM NaCl, 5% glycerol, 1 mM β-ME. A linear salt gradient from 0% to 40% IEX buffer B (20 mM Tris-HCl pH 8.0, 500 mM NaCl, 5% glycerol, 1 mM β-ME) was conducted. Proteins usually eluted in buffer containing 200 mM NaCl and fractions collected were analysed by SDS-PAGE.

### Isothermal titration calorimetry (ITC)

Human SNX1 and SNX2 peptide sequences were synthesized by Genscript (USA). To obtain the peptide stock solutions at neutral pH, 100 mM Tris-HCl, pH 10.6 was used to dissolve the peptide to a final concentration of 8 mM. The peptides were: SNX1_35-51_ EAGDSDTEGEDIFTGAA, SNX_135-51(D45K)_ EAGDSDTEGEKIFTGAA, SNX1_75-92_ NGIHEEQDQEPQDLFADA, SNX2_16-33_ TDFEDLEDGEDLFTSTVS, SNX2_16-33(DLF/SSS)_ TDFEDLEDGESSSTSTVS, and SNX2_62-82_ TEVVLDDDREDLFAEATEEVS.

All microcalorimetry experiments were conducted at 25℃ using a PEAQ^TM^ ITC (Malvern) in 50 mM HEPES (pH 7.5), 200 mM NaCl, 5% glycerol and 1 mM β-ME (ITC buffer). All proteins were purified by SEC using the same buffer prior to ITC experiments to avoid buffer mismatches. To measure binding of SNX27 to SNX1 proteins, 800 μM of hSNX27_FERM_ was titrated into 20 μM of GST-mSNX1_1-139_ or His-SNX1_FL_. For the binding of SNX1 and SNX2 peptides to SNX27, peptides were diluted 10-fold to 800 μM in ITC buffer peptides and titrated into 20 μM SNX27_FL_ proteins containing the same 1:10 ratio of peptide buffer (100 mM Tris-HCl, pH10.6). The competitive ITC experiments were carried out by titrating 800 μM SNX1_75-92_ peptides into 20 μM proteins with 2-fold molar excess SNX2_16-33_ peptide pre-incubated with SNX27 on ice for an hour. For ITC experiment using SNX1_75-92_ against hSNX27 wild-type and single-site mutants, 600 μM SNX1_75-92_ peptide was titrated into 20 μM hSNX27 proteins.

The ITC experiments were conducted with one injection of 0.4 μL followed by a series of 12 injections of 3.2 μL each with 180 s intervals and a stirring speed of 850 rpm. The heat exchange of interactions was obtained through integrating the observed peaks and background correction by subtracting the heats of dilution. Data were analysed with Malvern software package by fitting and normalised data to a single-site binding model, yielding the thermodynamic parameters *K_d_*, *ΔH*, *ΔG*, and *-TΔS* for all binding experiments. The stoichiometry was refined initially, and if the value was close to 1, then N was set to exactly 1.0 for calculation. All experiments were performed at least twice to guarantee reproducibility of the data.

### AlphaFold predictions of peptide binding to SNX27

To generate predicted models of SNX1 and SNX2 N-terminal peptides associating with the SNX27 FERM domain we used the AlphaFold2 neural-network (Jumper et al., 2021) implemented within the freely accessible ColabFold pipeline (Mirdita et al., 2021). For each peptide-modelling experiment, the sequence of the human SNX27a FERM domain (residues 269 to 531; Q96L92) was modelled with the four separate N-terminal aDLF sequences from SNX1 and SNX2. The specific peptides we modelled were residues 33 to 56 and 75 to 98 from human SNX1 (Q13596), and residues 14 to 37 and 60 to 83 from human SNX2 (O60749). ColabFold was executed using default settings where multiple sequence alignments were generated with MMseqs2 (Mirdita et al., 2019) and structural relaxation of final peptide geometry was performed with Amber (Hornak et al., 2006) to generate three models per peptide. Sequence conservation was mapped onto the modelled SNX27 FERM domain structure with Consurf (Ashkenazy et al., 2016). Structural alignments and images were generated with Pymol (Schrodinger, USA).

### Phylogenetic analyses

Homologous proteins were identified using NCBI protein-protein BLAST (with default parameters) against all non-redundant GenBank CDS Translations+PDB+SwissProt+PIR+PRF excluding environmental samples from WGS projects for a wider range of species (unaligned sequence data with accession numbers available in supplemental material). For the ctenophore data we used the BLAST (with default parameters) from the Mnemiopsis Genome Project Portal (MGP) portal (Ryan *et al*. 2013, Moreland *et al*. 2014, Moreland *et al*. 2020). Sequences were aligned using MAFFT v7.480 (Katoh and Standley 2013), L-INS-I mode, and then used to infer initial exploratory trees to identify specific duplications (using IQtree 2.1.4 (Minh *et al*. 2020) with the LG+F+G model and 1000 ultrafast bootstraps. After orthologues of each copy of the gene were identified in a range of taxa, we then performed iterative tree inferences to confirm paralogues and to identify species-specific duplication events. Subsequent maximum likelihood phylogenetic analyses were performed using IQtree 2.1.4 with the best fitting model selected using the Bayesian information criterion (BIC), including models which account for site-rate heterogeneity (LG+C10…C60) (Si Quang et al., 2008), empirical amino acid frequencies (+F), and accounting for across-site rate variation using either a Gamma distribution (+G) (Yang, 1994) or the free rates model (+R) (Yang, 1995; Soubrier et al., 2012) with 10000 ultrafast bootstrap replicates. The best fitting model according to BIC was LG+C30+F+G for the final SNX1 and SNX2 tree (see Supplemental Material), and LG+C20+F+G for SNX27 and VPS26 (alignments and maximum likelihood trees in Supplemental Material).

## ACKNOWLEDGEMENTS

We thank the Wolfson Bioimaging Facility at the University of Bristol for their support and Dr Da Jia for discussion. Work in the Cullen laboratory is supported by the Wellcome Trust (104568/Z/14/Z and 220260/Z/20/Z), the Medical Research Council (MR/L007363/1 and MR/P018807/1), the Lister Institute of Preventive Medicine, and the award of a Royal Society Noreen Murray Research Professorship to P.J.C. (RSRP/R1/211004). B.M.C. is supported by a Senior Research Fellowship and Project Grant from the National Health and Medical Research Council (APP1136021 and APP1156493). E.R.R.M. is supported by a Research Fellows Enhancement Award to T.A.W (RGF\EA\180199). T.A.W. is supported by a Royal Society University Fellowship (URF\R\201024).

## AUTHOR CONTRIBUTIONS

Biochemistry and cell biology analysis: B.S., M.G-A. and C.M.D. Structural analysis and ITC: Q.G. and K.C. Evolutionary analysis: E.R.R.M. and T.A.W. Isolation of SNX27 KO line: A.J.E. Data analysis and Figure generation: B.S., Q.G., M.G-A., K.C. and E.R.R.M. Manuscript Writing - 1^st^ draft: B.S. and P.J.C; Final Version: all authors. Initial Concept: B.S., B.M.C. and P.J.C. Concept Development: all authors. Funding: T.A.W., B.M.C. and P.J.C. Supervision: B.S., T.A.W., B.M.C. and P.J.C.

