## Supplementary Data for "Mechanistic basis for SNX27-Retromer coupling to ESCPE-1 in promoting endosomal cargo recycling"

**Supplementary Figure 1**

**A** Co-immunoprecipitation of GFP-tagged extended N-terminal regions (NT) of SNX1 and SNX2 alongside the GFP-tagged short N-terminal regions of SNX5 and SNX6 expressed in HEK293T cells. The cell lysates were subjected to GFP-trap based immunoprecipitation and the immunoprecipitates were blotted for SNX27 and GFP. The blot is representative of three independent GFP traps.

**B** Competitive ITC assay of $SNX2_{16-33}$ and $mSNX27_{FL}$ pre-incubated with 2-fold molar excess of $SNX1_{75-92}$ peptide showing that binding is blocked by the competing $SNX1_{75-92}$ peptide.

Molecular masses are given in kilodaltons.

**Supplementary Figure 2**

As described in methods, twelve models of SNX27 were generated in total, consisting of three models each of SNX27 in association with the four aDLF sequences from human SNX1 and SNX2. The top panel shows the overlay of all twelve models in C$\alpha$ trace representation coloured by the pLDDT score (predicted Local Distance Difference Test). The pLDDT is a per-residue confidence score between 0 (lowest confidence) and 100 (highest confidence). The majority of the SNX27 structures show very high pLDDT scores and overlay with very high precision. The lower panel shows only the best eight models where the core aDLF sequences have identical binding conformations.

**Supplementary Figure 3**

Maximum likelihood phylogenies of (A) SNX27, (B) VPS26A/B and (C) VPS5/SNX1/SNX2.

**A** SNX27 (green) with SNX17 (red) used as an outgroup with representatives across Filozoa.

**B** VPS26A, VPS26B & VPS26C with VPS26C in red, and VPS26B in green, the duplication event occurring in the ancestral vertebrate is in blue.

**C** The SNX1 sequences across choanozoans, with the duplication in vertebrates in blue. Phylogenies were inferred under the best-fit model as determined by BIC. Branch support values are percentages based on 10000 ultrafast bootstraps.

#### Supplementary Table 1. Thermodynamic parameters for the binding of SNX1/SNX2 with SNX27 by ITC.

| Syringe | Cell | $K_d$ (μM) | *ΔH* (kcal/mol) | *ΔG* (kcal/mol) | *-TΔS* (kcal/mol) |
|---|---|---|---|---|---|
| **hSNX27$_{FERM}$ against mSNX1 proteins** | | | | | |
| hSNX27$_{FERM}$ | mSNX1$_{1-139}$ | 39.10 ± 0.42 | -6.93 ± 0.01 | -6.02 ± 0.01 | -0.92 ± 0.02 |
| hSNX27$_{FERM}$ | mSNX1$_{FL}$ | 41.00 ± 1.82 | -6.23 ± 3.11 | -5.99 ± 0.02 | 0.24 ± 3.10 |
| **hSNX1/2 peptides against mSNX27$_{FL}$** | | | | | |
| hSNX1$_{35-51}$ | mSNX27$_{FL}$ | 33.43 ± 1.89 | -1.72 ± 0.65 | -6.11 ± 0.04 | -4.39 ± 0.64 |
| hSNX1$_{35-51}$ (D45K) | mSNX27$_{FL}$ | No binding detected | | | |
| hSNX1$_{75-92}$ | mSNX27$_{FL}$ | 13.37 ± 0.75 | -1.98 ± 0.18 | -6.65 ± 0.03 | -4.68 ± 0.17 |
| hSNX2$_{16-33}$ | mSNX27$_{FL}$ | 37.8 ± 1.44 | -0.85 ± 0.30 | -6.04 ± 0.02 | -5.19 ± 0.29 |
| hSNX2$_{16-33}$ (DLF-SSS) | mSNX27$_{FL}$ | No binding detected | | | |
| hSNX2$_{62-82}$ | mSNX27$_{FL}$ | 22.63 ± 1.89 | -1.85 ± 0.43 | -6.34 ± 0.05 | -4.49 ± 0.43 |
| **Competitive ITC assay of hSNX2$_{16-33}$ and hSNX1$_{75-92}$ with mSNX27$_{FL}$** | | | | | |
| hSNX2$_{16-33}$ | mSNX27$_{FL}$ & hSNX1$_{75-92}$ | No binding detected | | | |
| hSNX1$_{75-92}$ | mSNX27$_{FL}$ & hSNX2$_{16-33}$ | 30.35 ± 1.06 | -2.56 ± 0.14 | -6.17 ± 0.02 | -3.61 ± 0.12 |
| **hSNX1 peptide 75-92 against hSNX27$_{FL}$ WT and mutants** | | | | | |
| hSNX1$_{75-92}$ | hSNX27$_{FL}$ | 12.77 ± 0.47 | -1.65 ± 0.54 | -6.68 ± 0.03 | -5.03 ± 0.54 |
| hSNX1$_{75-92}$ | hSNX27$_{FL}$ K495D | No binding detected | | | |
| hSNX1$_{75-92}$ | hSNX27$_{FL}$ K496D | 56.40 ± 2.83 | -1.30 ± 1.07 | -5.80 ± 0.03 | -4.50 ± 1.04 |
| hSNX1$_{75-92}$ | hSNX27$_{FL}$ R498D | No binding detected | | | |
| hSNX1$_{75-92}$ | hSNX27$_{FL}$ K501D | No binding detected | | | |

### *Supplementary Figure 1*

*A*

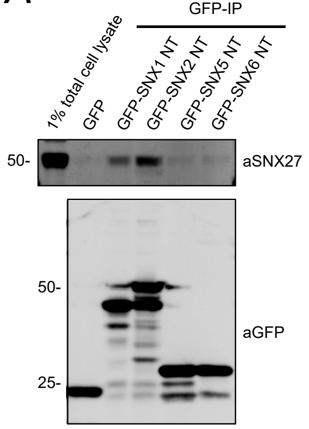


*B*

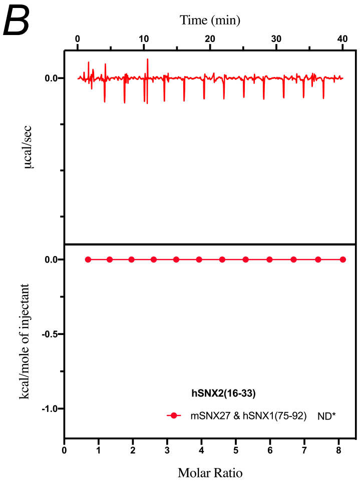

### Supplementary Figure 2

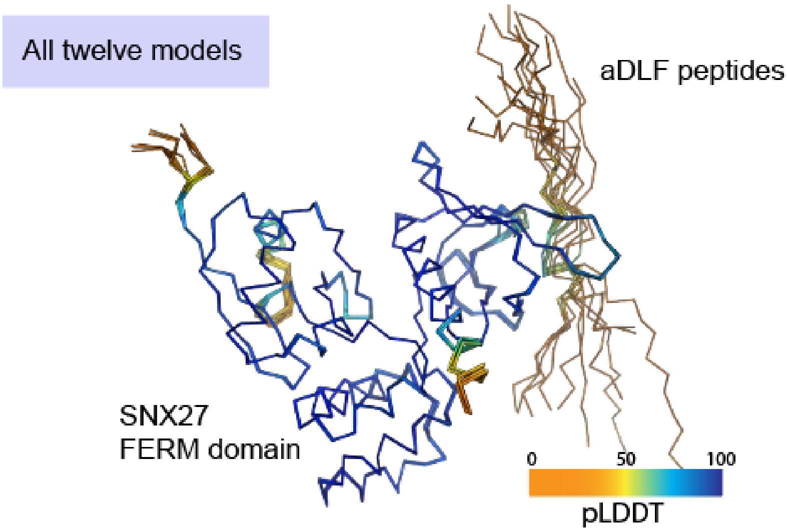


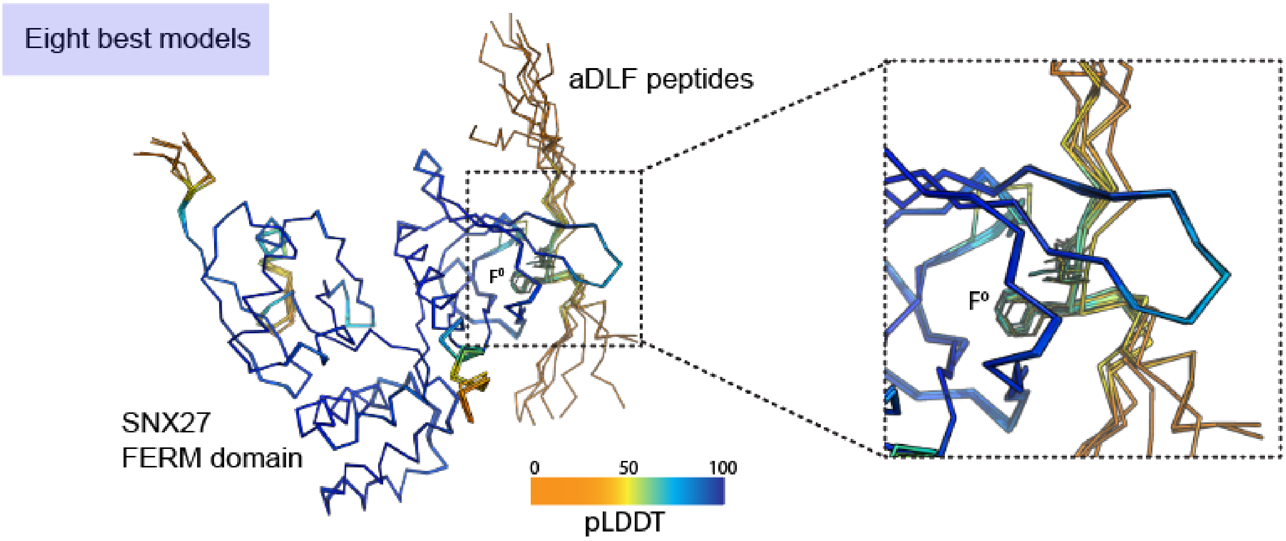

### Supplementary Figure 3

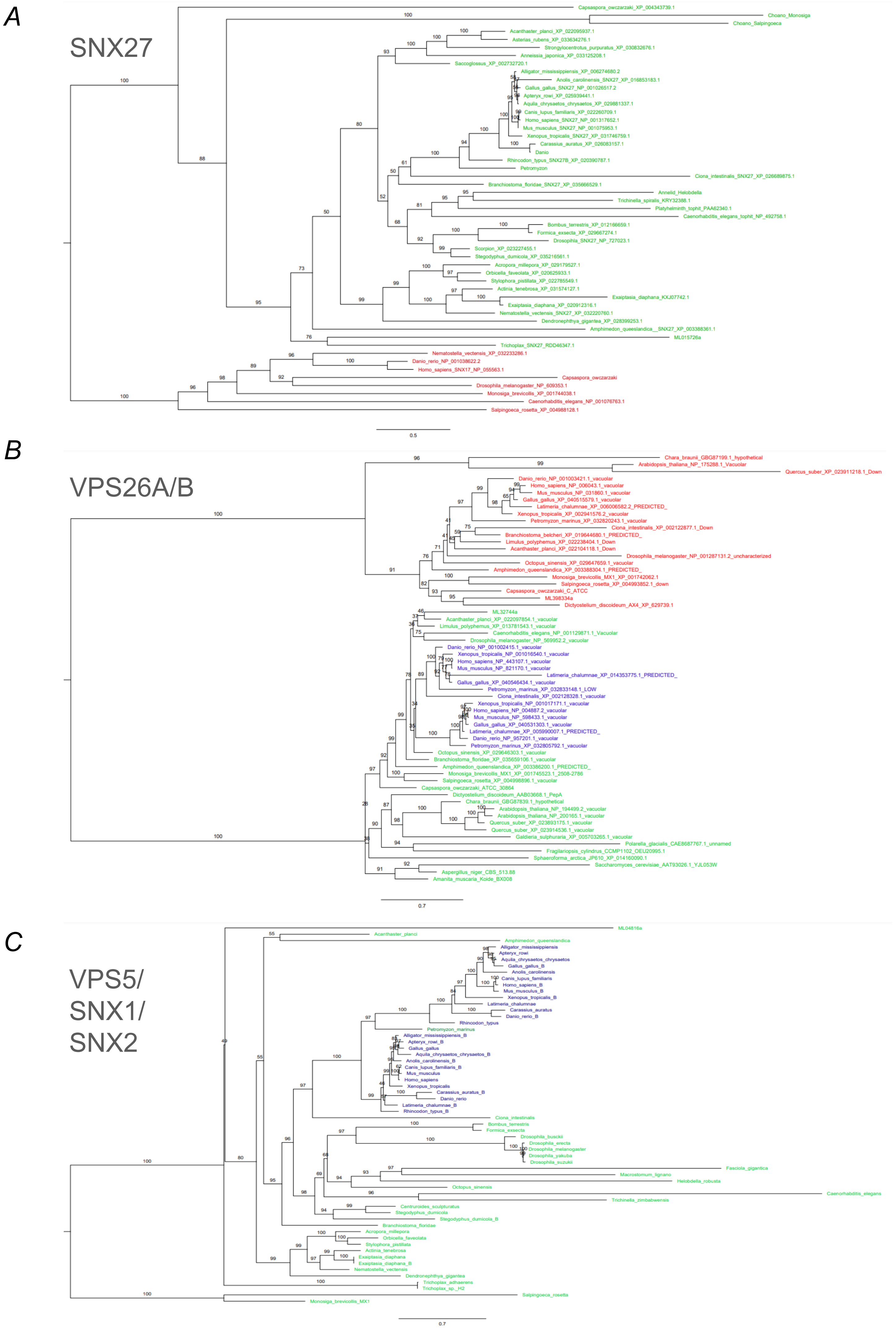
